# Peptides alleviate cognitive impairment by inhibiting and disassembling amyloid-β aggregates in Alzheimer’s disease

**DOI:** 10.1101/2025.05.28.656730

**Authors:** Wenqing Gao, Baocai Gao, Sen Wang, Di Wu, Pucheng Mei, Lu Geng, Chuhao Li, Yuanyuan Li, Xuehe Liu, Jiasong Pan, Jing Huang, Yifan Deng, Xiangjun Chen, Hexige Saiyin, Hongbo Yu, Suhua Li, Qiuxin Zhang, Bing Xu, Fang Huang, Chen Ming, Hao Wu, Jixi Li

## Abstract

Alzheimer’s disease (AD) is a devasting neurodegenerative disorder characterized by β-amyloid formation, further exacerbated by RIPK1/RIPK3 necrosome-induced programmed necrosis (necroptosis). We previously showed that the RIPK1/RIPK3 necrosome forms a functional amyloid complex using its RIP homotypic interaction motifs (RHIMs). Here, we discovered that the core RIPK1/RIPK3 necrosome shares strikingly structural similarity to the C-terminal region of β-amyloid (Aβ42), and the RHIM-derived tetrapeptides (IQIG or VQVG) directly inhibit Aβ aggregation, disassemble preformed Aβ fibrils (PFFs), and reduce RIPK1 polymerization. Also, the peptides exhibit effective membrane permeability and could reduce Aβ40 or Aβ42-induced neural death and TNFα-induced necroptosis in SH-SY5Y cells. IQIG and VQVG injected by ICV increase learning and memory abilities by reducing Aβ plaques and hyperphosphorylated tau in the cortex and hippocampus of APP/PS1 double-transgenic mice. Mechanistically, the peptides directly interact with Aβ to block Aβ aggregation and alleviate microglia-mediated neuroinflammation. Strikingly, single-cell RNA sequencing revealed that the peptides-treated AD transgenic mice restored neuronal homeostasis with increased GABAergic neurons and decreased glutamatergic neurons. Furthermore, total cell-cell interaction strength increased while the AD risk gene Apoe expression decreased in the specific oligodendrocyte subtype of peptides-treated AD mice. Thus, our findings revealed that the peptides could improve cognition and memory capabilities and serve as promising structural templates for potential drugs against AD.

## INTRODUCTION

Alzheimer’s disease (AD) is the most prevalent type of dementia, affecting approximately 45 million people worldwide. Neural cell death, especially in the cortex and hippocampus, is responsible for the decline of learning and memory abilities observed in AD patients ^1^. The histopathological hallmarks of AD consist of neurofibrillary tangles, caused by abnormally phosphorylated tau and amyloid plaques^2^. Thus far, the most widely accepted hypothesis for AD pathogenesis has focused on the gradual accumulation of β-amyloid (Aβ) ^3^. Individual Aβ40 and Aβ42 monomers can form different aggregates through amyloid-like interactions, soluble oligomers of different molecular weights, fibrils, and large plaques, to mediate neurodegeneration in AD ^4^. Furthermore, disaggregation of Aβ oligomers can decrease Aβ-induced neuroinflammation and rescues cognitive deficits in APP/PS1 double-transgenic mice^5^. Therefore, Aβ aggregation or protofibrils are believed to represent critical structures that produce cytotoxicity, contribute to synaptic deficits, and initiate the detrimental cascade involved in the pathology of AD ^6–8^. Recently, the anti-Alzheimer’s drugs Lecanemab and Donanemab gained traditional FDA approval, indicating that targeting soluble Aβ oligomers or protofibrils by antibody is a promising treatment for AD ^8, 9^. Moreover, structural-based small peptides have been used as inhibitors of amyloid or Tau aggregation in vitro ^10, 11^.

Increasing evidence suggests that necroptotic cell death is highly associated with AD progression, with increased necroptosis markers in brains of 11-month-old 5xFAD (Familial Alzheimer’s disease) transgenic mice overexpressing mutant human Aβ and postmortem human AD brains ^12–14^. Receptor-interacting protein kinase 1 (RIPK1, or RIP1) and RIPK3 (or RIP3) form a large complex called necrosome, which plays critical roles in necroptosis induced by multiple stimuli ^15, 16^. The anti-necroptotic molecule necrostatin-1 (Nec-1), which inhibits the RIPK1 kinase activity explicitly, can reduce neuroinflammation mediated by microglia, making RIPK1 a promising therapeutic target for the treatment of AD ^17–20^.

RIPK1 and RIPK3 contain a Ser/Thr kinase domain (KD) and a RIP homotypic interaction motif (RHIM). In addition, RIPK1 also has a death domain (DD) at its C terminus for recruitment to the TNF receptor signaling complex ^21, 22^ (Fig. 1a). We previously showed RIPK1 and RIPK3 assemble into a hetero-oligomeric functional amyloid complex via the RHIMs ^23^. The RIPK1/RIPK3 hetero-amyloid consists of a pair of meandering, parallel β-sheets broken into four short β-segments by short turns; the two sheets come together in an antiparallel fashion to create a hydrophobic core ^23, 24^. The critical interactions in the core involve residues from each chain (532-549 in RIPK1 and 448-462 in RIPK3), near their IQIG (RIPK1) and VQVG (RIPK3) consensus sequence elements ^24^ (Fig. 1a and S1a).

**Figure 1.**
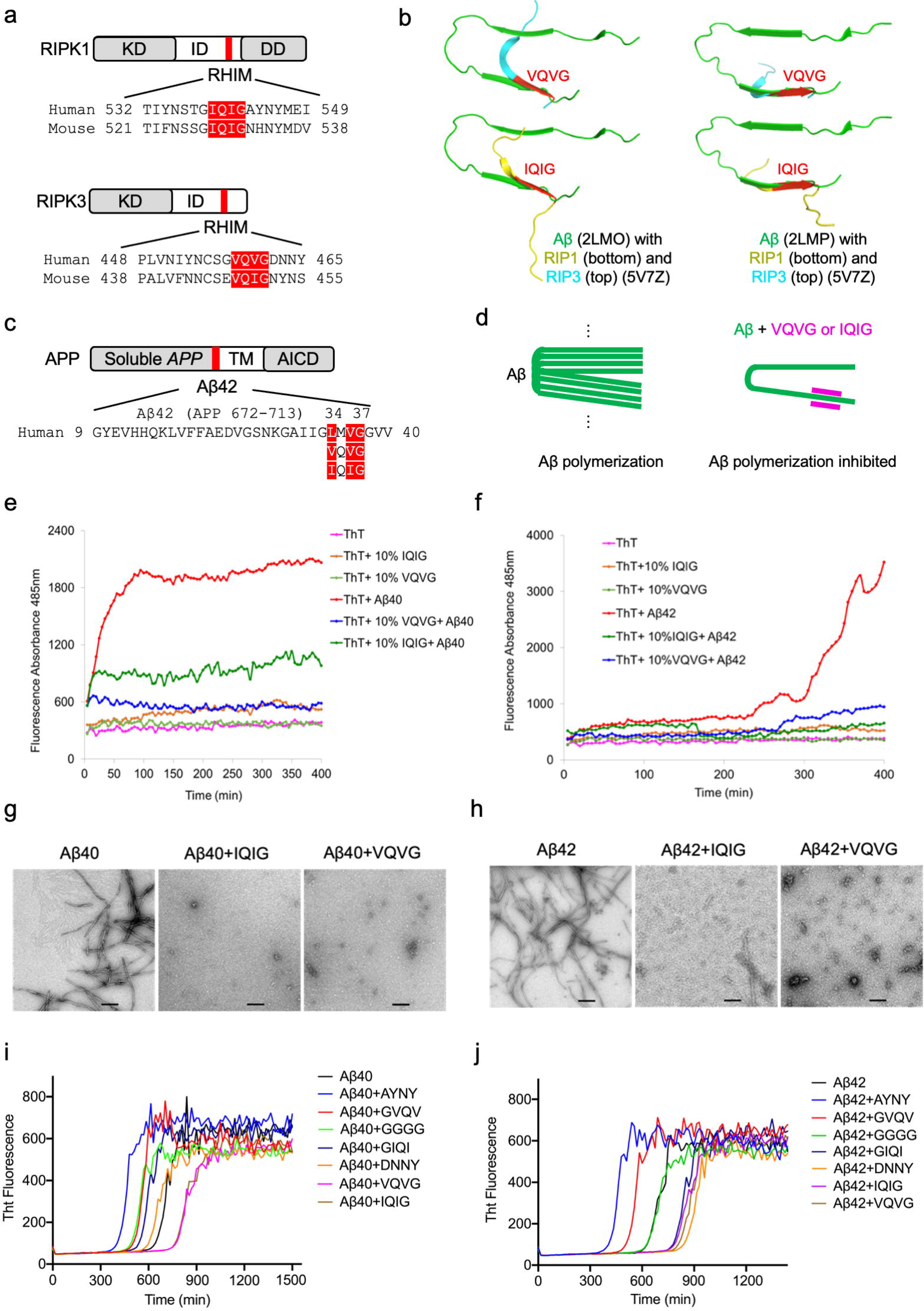
IQIG and VQVG inhibited Aβ aggregation in vitro. **a**. Domain structures of RIPK1 and RIPK3 and the alignment of RHIMs. Length is indicated in the number of amino acids. KD, kinase domain; ID, intermediate domain; RHIM, RIP homotypic interaction motif; DD, death domain. The RHIM consensus IQIG and VQV(I)G from human and mouse RIPKs are shown in red. **b**. Unrestrained protein structure alignment of Aβ amyloid (PDB 2LMO and PDB 2LMP, green), RIPK1 (PDB 5V7Z, yellow), and RIPK3 (PDB 5V7Z, blue). **c**. Domain structure of the amyloid precursor protein (APP) and amino acid sequence alignment of Aβ42, IQIG, and VQVG. Identical sequences are shaded in red. Length is indicated in the number of amino acids. TM, transmembrane. AICD, the APP intracellular domain. Aβ42 domain is highlighted in red. **d**. Schematic representation for the potential working mechanism of IQIG or VQVG (purple) on its inhibition of Aβ polymerization (green). **e-f**. The polymerization of Aβ40 (**e**) or Aβ42 (**f**) was monitored by the thioflavin T (ThT) fluorescence. Aβ (10 μM) was incubated with 20 μM ThT and the corresponding ratio of IQIG or VQVG peptides at 25 °C. **g-h**. Analysis of Aβ40 or Aβ42 aggregates by negative staining electron microscopy. Aβ40 (20 μM) and Aβ42 (20 μM) were incubated with or without the tetrapeptide IQIG (2 μM) or VQVG (2 μM) for 24 h. Scale bars, 100 nm. **i-j**. The polymerization of Aβ40 (**i**) or Aβ42 (**j**) was monitored by the ThT fluorescence. Aβ (10 μM) was incubated with 20 μM ThT and the corresponding ratio of IQIG, VQVG, GIQI, GVQV, AYNY, DNNY, and GGGG peptides at 25 °C. All data are representative of three independent experiments.

Here, we incidentally discovered structural and sequence similarity between Aβ and the necrosome at their core sequences and found that the short RHIM peptides, IQIG and VQVG, inhibited the oligomerization of both Aβ and RIPKs in vitro. When used in cells, they decreased necroptosis, exhibited protective effects on Aβ aggregates-induced neuronal cell death, and reduced the levels of Aβ oligomers and fibrils. Furthermore, IQIG and VQVG could reduce Aβ plaques and hyperphosphorylated tau, and provide cognitive benefits for amyloid-rich aged double transgenic APP/PS1 mice expressing a chimeric mouse/human amyloid precursor protein (APP) and a mutant human presenilin 1 (PS1). Thus, the IQIG and VQVG peptides may directly inhibit Aβ aggregation and necroptosis from serving as promising structural templates for potential drugs against AD.

## METHODS

### Animals

APP/PS1 transgenic mice that overexpress mouse/human amyloid precursor protein (Mo/Hu APP695 SWE) and mutant human presenilin-1 (PS1-△9) in a C57BL/6J genetic background were purchased from Shanghai Model Organism Ltd (China, Shanghai; Animal certificate number: 20170010000109). The mouse genotype was confirmed by PCR using DNA extracted from tail tissues. All mice were housed under standard conditions at 22 °C with a 12-h light/dark cycle and free access to food and water. Animal care and experimental protocols used in this study were conducted according to the guidelines approved by the Care and Use of Laboratory Animals of Fudan University. All *in vivo* experiments were done in a randomized and blinded fashion. Effects were made to minimize animal suffering and to reduce the number of animals used.

### Reagents and antibodies

Peptides Aβ40 and Aβ42 used for cell viability and ThT assays were synthesized from Sangon Biotech Co., Ltd (Shanghai). The tetrapeptide IQIG, VQVG, GIQI, GVQV, AYNY, DNNY, GGGG, LMVG, and FITC-labeled peptides were synthesized from ChinaPeptides Co., Ltd (Shanghai). The IQIG-NBD, VQVG-NBD, and bb-NBD were synthesized according the methods in previous study ^25^. TNF-α, Smac mimetic, and Z-VAD were purchased from Sigma-Aldrich. The antibodies used for immunoblotting were anti-Aβ (1:5000, ab2539, Abcam), anti-oligomer A11 (1:1000, AHB0052, Thermo Fisher), anti-RIPK3 (1:500, sc-374639, Santa Cruz Biotechnology), anti-RIPK1 (1:500, sc-133102, Santa Cruz Biotechnology), anti-β-actin (1:5000, Abcam) anti-MLKL (1:1000, ab243142, Abcam), and anti-p-MLKL (1:1000, ab196436, Abcam). The primary antibodies for immunostaining were as follows: rabbit anti-Aβ (1:100, ab20106, Abcam), rabbit anti-GFAP (1:100, ab68428, Abcam), rabbit anti-Iba1 (1:100, ab178846, Abcam), rabbit anti-p-tau (1:100, ab92676, Abcam), mouse anti-RIPK1 (1:100, sc-133102, Santa Cruz Biotechnology) and mouse anti-RIPK3 (1:100, sc-374639, Santa Cruz Biotechnology). Fluorescent secondary antibodies were purchased from Invitrogen (USA) and were used in a 1:200 dilution.

### Cell culture and treatment

HeLa, HEK293T, L929, and SH-SY5Y cells were grown in DMEM medium (Hyclone) supplemented with 10% fetal bovine serum (Gibco) at 37 °C with 5% CO_2_ in a humidified atmosphere. SY5Y cells were plated into a 96-well plate before treatment of Aβ aggregates, IQIG, or VQVG peptides. Aβ42 peptide was dissolved in DMSO as 10 mM stock, then diluted into 10-fold by cell starvation medium (0.5% fetal bovine serum in DMEM) and incubated for 24 h in 37 °C to polymerize different types of Aβ. Pre-aggregated Aβ was applied to the cells for 24 and 36 h, respectively, at a concentration of 10 μM. IQIG or VQVG (10 μM and 50 μM) was applied to the medium 15 min before applying Aβ aggregates.

### Plasmids

Aβ40 (residues 672-711 in human APP gene) or Aβ42 (residues 672-713 in human APP gene) were subcloned from the human APP cDNA (Uniprot ID: P05067) into the pEGFP-N1vector and pmCherry-N1, respectively. The human RIPK1 (residues 520-560 or 496-583) and human RIPK3 (residues 428-498) were subcloned from the full-length RIPK1 and RIPK3 cDNAs into the pmCherry-N1 vector, respectively ^23^. All constructs were confirmed by DNA sequencing.

### Protein expression and purification

The truncated RHIMs of human RIPK1(residues 520-560, or 496-583) and human RIPK3 (residues 428-498) were synthesized with a codon-optimization strategy in the pSMT3 vector with an N-terminal His-sumo tag (Genewiz, China) and were expressed in BL21-CodonPlus (DE3)-RIL cells. Briefly, cells were suspended in buffer A (20 mM Tris-HCl, pH8.0, 300 mM NaCl, 5% glycerol, 0.2 mM DTT) and lysed by a high-pressure homogenizer. The RIPK1 and RIPK3 proteins were then purified by Ni-NTA affinity chromatography, and the His-sumo tag was removed by ULP1 cleavage followed by additional Ni-NTA affinity chromatography. The target proteins were then applied to a Superdex 200 gel filtration column pre-equilibrated in buffer B (20 mM Tris-HCl, pH7.4, 100 mM NaCl). The proteins were finally concentrated to around 10 mg ml^-1^. Aβ40 (residues 672-711 in human APP gene) or Aβ42 (residues 672-713 in human APP gene) was expressed and purified using the same method as RIPK1 and RIPK3.

### Cell viability assay

Cell viability was analyzed using an Enhanced Cell Counting Kit-8 Assay Kit (Beyotime, China) according to the manufacturer’s instructions. Briefly, 5×10^3^ cells were seeded in 96-well plates. After 12 h, the cells were treated with reagents for the indicated durations. The medium was then changed to a medium containing 5% CCK8-reagent (Beyotime, China) after 24 h and 36 h. The incubation was continued for one hour in the dark. Then, the absorbance was measured at a wavelength of 450 nm using a Spectra-Max M5 plate reader (Molecular Devices).

### Transmission electron microscopy (TEM)

The samples Aβ40, Aβ42, IQIG, or VQVG were negatively stained with uranyl formate and imaged with a Veleeta 2K x 2K cooled CCD camera (Olympus-Soft Imaging Solutions, Germany) on a JEM-1400 (JEOL, Japan) electron microscope at 120 kV. In brief, Aβ40 (20 μM) and Aβ42 (20 μM) proteins were incubated with or without the tetrapeptide IQIG (2 μM) or VQVG (2 μM) for 24 h, and then were observed by using negative staining TEM. For the disassembling assay, Aβ40 and Aβ42 preformed fibrils (PFFs, 20 μM) were incubated with or without IQIG 10 (μM) or VQVG (10 μM) peptides. The samples were observed by TEM during 12-32 h incubation.

### Blotting analyses and quantification of A***β*** in brain lysates

Brains of wild type mice or APP/PS1 mice and cells were lysed on the ice using a RIPA lysate supplemented with protease inhibitor cocktail (Roche) for 15 min and then centrifuged at 13,000 rpm for 15 min at 4 °C. The lysates were subjected to the SDS-PAGE analysis, transferred to nitrocellulose membranes, and blocked with 5% skim milk. After blocking, the membranes were incubated with primary antibodies overnight at 4 °C. Blots were washed with a TBST buffer and incubated with the horseradish peroxidase-conjugated secondary antibodies at room temperature for one hour before visualization with the ECL. Bands intensities were quantified using the Image J software program.

### Immunohistochemistry and immunofluorescence

The frozen sections of mouse tissues were warmed for 30 minutes at 37 °C before being blocked with 5% goat serum (Gibco) blocking solution for 30 min at 37 °C. Next, sections were incubated overnight at 4 °C with primary antibodies. After rinsing three times with PBS, sections were incubated with species-specific secondary antibodies and were visualized under fluorescence microscopy (Lecia TCS SP8, Germany). For immunofluorescence analysis, treated or untreated cells were fixed in 4% paraformaldehyde for 10 min at room temperature. The fixed cells were permeabilized with a blocking solution, including 0.1% Triton X-100 and 5% BSA in PBS for 30 min. Primary and secondary antibodies were diluted in PBS at the indicated dilutions and incubated for one hour at room temperature. Cells were washed in PBS, mounted in DAPI to counterstain the nucleus, and then observed under a fluorescence microscopy microscope (Zeiss-LSM880, Germany).

### Morris water maze

We used the standard water maze platform equipment for the repeated parallel experiments. The path length between the point of origin and the platform is 70-80 cm. A platform was placed 1.5 cm below the water surface in the second quadrant of a circular water tank (150 cm in diameter, 25 ± 2 °C). Water was made cloudy by adding titanium dioxide. On the first day, the mice were placed in a water tank and allowed to swim freely for 120 seconds. Mice were trained for 5 consecutive days with 4 trials per day as acquisition. For these trails, animals were placed at different starting positions spaced equally around the perimeter of the pool in random order. Animals were allowed to swim freely for 60 s to find the platform. If an animal could not locate the platform, it was guided to the platform and was allowed to stay on the platform for 20 s. The time that each mouse took to reach the platform was regarded as escape latency. The total distance that animals traveled in the target region was recorded. On the 7th day, a probe test was performed without the platform for 60 s. Each mouse was placed in the pool with the platform removed. The time they spent in the target zone and the number of crossings in the probe was recorded.

### Thioflavin T fluorescence

Thioflavin T (ThT, Sigma) was dissolved in DMSO to obtain a 0.1CM stock solution. ThT, Aβ40, Aβ42, IQIG, and VQVG stock solutions were diluted with the PBS buffer. Aβ peptides (30 μM) were centrifuged at 17,000×g for 20 min, and then the supernatant was retained for subsequent experiments. These reagents were mixed at a ratio of 1:1:1 so that the final concentration of Aβ was 10 μM. All mixtures were added into a black-walled 96-well plate and incubated at 25 °C, and the fluorescence signals were detected with a time-course on a Spectra-Max M5 plate reader (Molecular Devices). The excitation wavelength was 430 nm, and the emission wavelength was 485Cnm.

### ThS staining of A***β*** plaques in brain sections

The excised brains of wild type mice or APP/PS1 mice were fixed in 4% paraformaldehyde (pH 7.4) for 16 hours. The frozen brain samples were then immersed in 30% sucrose for cryoprotection and cut into 40 μm thick slices using a Cryostat (Leica Biosystems, Germany). To visualize the Aβ plaques, the sliced brains were stained with thioflavin S (ThS, Sigma) for 7 minutes. 500 μM of ThS was dissolved in 50% ethanol. After rinsing with 100%, 95%, and 70% ethanol successively, the sections were moved into PBS buffer. Images were taken with a fluorescence microscopy microscope (Zeiss-LSM880, Germany). The plaque number was calculated from a single brain image of each mouse by using the Image J software program.

### ELISA

The pro-inflammatory response was determined in hippocampal lysates using the Cytokine & Chemokine 36-Plex Mouse ProcartaPlex™ Panel 1A for six cytokines (IFN-γ, IL-4, IL-10, IL-13, TNF-α, and MCP-1) following the protocol provided by the supplier (ThermoFisher, USA). Briefly, 50 μl of the diluted sample, calibrator, or control was added per well. The plate was sealed with an adhesive plate seal and incubated at room temperature with shaking for 2 h. Then, the plate was washed three times and the detection antibody was added with shaking for 2 h. Finally, the plate was washed, and the read buffer was added. Signals were measured on a SECTOR Imager 2400 reader (MSD, USA).

### Tissue extraction and library preparation for the scRNA-seq data

The single-cell RNA sequencing procedure was done on the 10X Chromium platform. There were 10,111 cells sequenced at first. We did the quality control following the standard pre-processing workflow of Seurat v4.3.0 ^26^. Cells with more than 20% of mitochondrial counts, or with less than 200 or more than 6,000 unique genes detected were excluded. Doublets were filtered based on DoubletFinder. Cells with multiple cell type markers were also excluded as possible doublets. After quality control, 8,934 cells (88.4 %) were left for afterward analysis. The scale factor in the normalization step was set as 10,000. The top 2,000 highly variable features were used for afterward clustering. The clustering was done by the FindClusters function of Seurat, where the resolution is set as 0.5. There were 36 clusters found at first. We further excluded small clusters with multiple cell-type components. There were 30 clusters left for afterward analysis.

The cell type annotation was done in two steps. In the manual curation step, the classical marker genes of major cell types were used. We further projected our scRNA-seq dataset onto the mouse Whole Cortex and Hippocampus 10X reference UMAP structure from the Allen Brain Atlases ^27^, where we only used the hippocampus part as a reference. Each cluster’s projection score and cellular component estimation were calculated using the MapQuery method of Seurat. Cell type proportion test was based on scProportionTest ^28^. Differential expression analysis was done for each cell cluster between two contrasts (i.e., IQIG-treated vs. control mice, and VQVG-treated vs. control group), based on the non-parametric Wilcoxon rank sum test through the FindMarkers function of Seurat. Cell-cell communication analysis was done using CellChat.

### Statistical analysis

Data were analyzed using Graphpad version 6 (Graphpad Software, La Jolla, CA) and presented by mean ± SD. Differences among the groups were examined using the one- or two-way ANOVA test. Results were considered to be statistically significant if *P < 0.05*.

## RESULTS

### The core RHIM peptides IQIG and VQVG directly inhibit Aβ aggregation

Given that Aβ and RHIM sequences all form amyloidal structures with stacked β-strands in their core structures ^24, 29–33^ (Fig. S1a), we analyzed their potential similarity by structural alignment. We used two Aβ amyloid structures derived from solid-state nuclear magnetic resonance (NMR), a dimer (PDB 2LMO) and a trimer (PDB 2LMP), and one Aβ40 structure derived from cortical tissue of an AD patient with the electron microscopy method (PDB 6W0O), with comparable β-hairpins as the building block for the stacked β-sheet ^29–32^ (Fig. S1b-S1c). The corresponding RMSD values were 0.473Å, 0.326Å, and 0.216Å, respectively. Furthermore, the unrestrained structural alignment showed that the RIPK1 structure (from PDB 5V7Z)^24^ aligned with the β-hairpin in both Aβ structures right at the IQIG sequence (Fig. 1a-1b). Analogously, the RIPK3 structure (from PDB 5V7Z) ^24^ aligned with the β-hairpin in both Aβ structures at the VQVG sequence despite divergent at other regions of the structures (Fig. 1a-1b). It turned out that the aligned region on the Aβ structures has the LMVG sequence, which bears similarity to the core RHIM sequence at 1^st^, 3^rd^ and 4^th^ positions and is, therefore, an RHIM-like sequence (Fig. 1c). Our aggregation experiments with thioflavin T (ThT), a specific dye for binding with β-strands in amyloid, revealed that mutating LMVG to AAAA or deleting LMVG from the entire sequence of Aβ produced the blocking effects on the aggregation progression (Fig. S1d-S1e). Similarly, the crystal structure of the VQVG peptide (PDB 5ZCK) ^24^ aligned with the three Aβ structures as well as the RIPK1 and RIPK3 peptide structures at the RHIM or RHIM-like consensus sequence (Fig. S1c). Because the RHIM peptides VQVG and IQIG do not form amyloid in solution, unlike their longer counterparts ^24^, we wondered if these peptides could inhibit the aggregation of Aβ as well as RIPK1 and RIPK3 by capping and stopping amyloid filament nucleation and growth (Fig. 1d).

To investigate whether RHIM-derived peptides of RIPK1 and RIPK3, IQIG and VQVG, could regulate Aβ aggregation, we mixed Aβ40 or Aβ42 monomers at 37 °C with the IQIG or VQVG peptide at a 10% molar ratio (Aβ: peptide = 10: 1). We then monitored Aβ40 amyloid formation using ThT, giving strong fluorescence upon binding to amyloids. Aβ40 or Aβ42 quickly polymerized by themsevles, while IQIG or VQVG treatment markedly inhibited the polymerization of Aβ40 or Aβ42, evidenced by reduced fluorescence signal (Fig. 1e-1f). To further confirm this conclusion, we used a negative-staining transmission electron microscope (TEM) to observe the morphologies of the samples after 24 h incubation. While both Aβ40 or Aβ42 alone samples formed entangled filaments, the IQIG- or VQVG-treated amyloid samples showed no filaments or only a few shorter filaments (Fig. 1g-1h). Next, we assayed five other tetrapeptide candidates to verify the specificity of IQIG and VQVG’s effect on Aβ aggregation. Of the five, four were also derived from RIPK1 or RIPK3 RHIM sequence, including AYNY (RIPK1_543-546_), GIQI (RIPK1_538-541_), DNNY (RIPK3_462-465_), and GVQV (RIPK3_457-460_) (Fig. 1a). The other one is the irregular control peptide GGGG ^34^. The results showed that IQIG and VQVG efficiently inhibited Aβ40 and Aβ42 aggregation, while GGGG, AYNY, and GVQV did not affect Aβ40 and Aβ42 aggregation (Fig. 1i-1j). Interstingly, DNNY and GIQI (represented the reversed sequence of IQIG) did not affect on Aβ40 aggregation, but had some inhibition abilities on Aβ42 amyloid formation (Fig. 1i-1j). Thus, these data revealed that RHIM tetrapeptides, IQIG and VQVG, could directly inhibit Aβ aggregation as suitable candidates.

### IQIG and VQVG disassemble preformed-fibrils (PFFs) of Aβ40 or Aβ42 and exhibited effective membrane permeability in cells

The potential therapeutic strategy to treat AD is to reduce amyloid plaques ^35^. However, few molecules can disintegrate preformed fibrils (PFFs) ^36–38^. We next investigate whether IQIG and VQVG could reverse the aggregation process and disassemble Aβ PFFs. The Aβ40 PFFs incubated with or without peptides were first monitored by the ThT fluorescence. The results showed that the ThT values decreased dramatically by 30-40% when adding IQIG or VQVG (PFFs: peptide=4:1) for 12 h incubation (Fig. 2a), indicating that the peptides could directly disaggregate Aβ40 fibrils. Next, we observed the morphologies of Aβ40 PFF samples with or without peptides treatments by the TEM method. It showed that the Aβ fibrils were reduced markedly and converted into granular aggregates after incubation with IQIG or VQVG for 12 h. In contrast, untreated Aβ40 fibrils kept the well-developed fibrillar morphology (Fig. 2b). This result indicated that IQIG and VQVG could disassemble Aβ40 PFF, correlating well with the ThT fluorescence result. Also, the addition of IQIG or VQVG to Aβ42 PFFs resulted in a sharp decrease of ThT fluorescence at 32 h (Fig. 2c). Further observation showed that the Aβ42 PFFs had been disaggregated into many shorter, thinner fibrils, or small spherical particles in the presence of peptides IQIG or VQVG (Fig. 2d). These results suggest that IQIG and VQVG are capable of dissociating preformed fibrils of Aβ40 and Aβ42.

**Figure 2.**
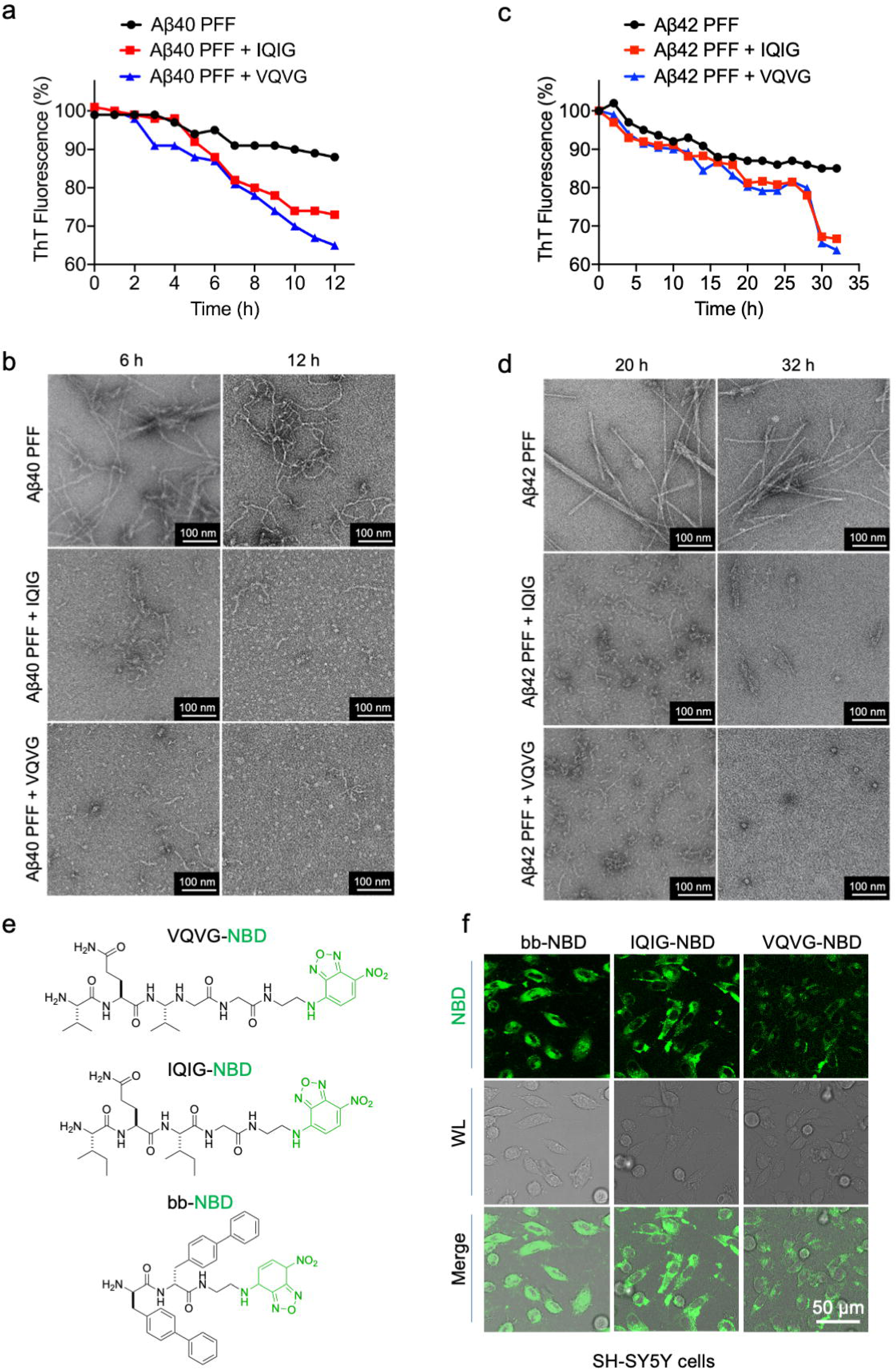
IQIG and VQVG disassembled Aβ preformed fibrils (PFFs) and exhibited effective membrane permeability. **a, c**. The disassembling effects of peptides on Aβ40 and Aβ42 PFFs were monitored by the ThT fluorescence. 20 μM Aβ40 (**a**) or Aβ42 (**c**) PFFs were incubated for 48 h, and then added 10 μM IQIG or VQVG peptides. The 485 nm absorbances for Aβ PFF itself (black) and with IQIG (red) or VQVG (blue) were shown. **b, d**. The Aβ40 (**b**) or Aβ42 (**d**) PFF samples treated with or without IQIG or VQVG were observed by TEM during 12-32 h incubation. The scale bar represents 100 nm. **e.** The chemical structrues of synthesized peptides VQVG-NBD, IQIG-NBD, and bb-NBD. The NBD group is shown in green. **f**. SH-SY5Y cells were treated with 2 μM bb-NBD, 5 μM IQIG-NBD, and 5 μM VQVG-NBD, respectively, for 24h before observation by the confocal microscopy. Scale bars, 50 μm.

To further investigate the cellular uptake of peptides in live cells, we conjugated NBD, a fluorophore (green), to the C-terminus of IQIG and VQVG (Fig. 2e). The peptide bb-NBD ^25^, comprising D-BiP (or b) and C-terminally capped with NBD, self-assembles into micelles. bb-NBD, used as a positive control, rapidly enters cells and localizes to membrane-rich organelles such as the ER, Golgi apparatus, and lysosomes ^25^. Confocal laser scanning microscopy (CLSM) was employed to visualize the uptake of IQIG-NBD and VQVG-NBD in SH-SY5Y and HeLa cells (Fig. 2f and S2c), showing significant intracellular enrichments upon treatment with peptides within 24 hours. Also, we examined the cell-membrane penetration ability using FITC-labeled tetrapeptides. SY5Y cells were transfected with Aβ42-mCherry for 24 h, and then treated with or without FITC-labeled tetrapeptides for 6 h. The peptides exhibited effective membrane permeability and could colocalize with Aβ42 puncta, indicating their potential function in inhibiting the formation of Aβ plaques (Fig. S2d).

### IQIG and VQVG inhibit Aβ-induced neural cell death

To investigate whether IQIG and VQVG could inhibit Aβ-induced cell death, we first examined whether they affected the formation of Aβ plaques in cells. Human neuroblast SH-SY5Y cells were transiently transfected with Aβ40-GFP or Aβ42-GFP, and the formation of Aβ plaques was assessed 24 h after treatment. We found that IQIG or VQVG treatments significantly downregulated Aβ plaque formation in the transfected SH-SY5Y cells (Fig. 3a-3b). Meanwhile, we prepared Aβ42-derived aggregates by incubating their monomers at 37 °C overnight, and then added them to SH-SY5Y cells using the CCK8 assay to assay the cell viability. The result showed that cell viability significantly decreased in all cultured cells treated with Aβ42 aggregates compared to non-treated cells at 24 h and 36 h time points (Fig. 3c). Also, when the Aβ aggregates-cultured SH-SY5Y cells were treated with IQIG or VQVG, the cell viability conspicuously increased, indicating that IQIG and VQVG effectively blocked the cytotoxicity induced by Aβ aggregates (Fig. 3c). Moreover, VQVG with a higher concentration maintained better neuroprotective effects at the more extended time point of 36 h (Fig. 3c).

**Figure 3.**
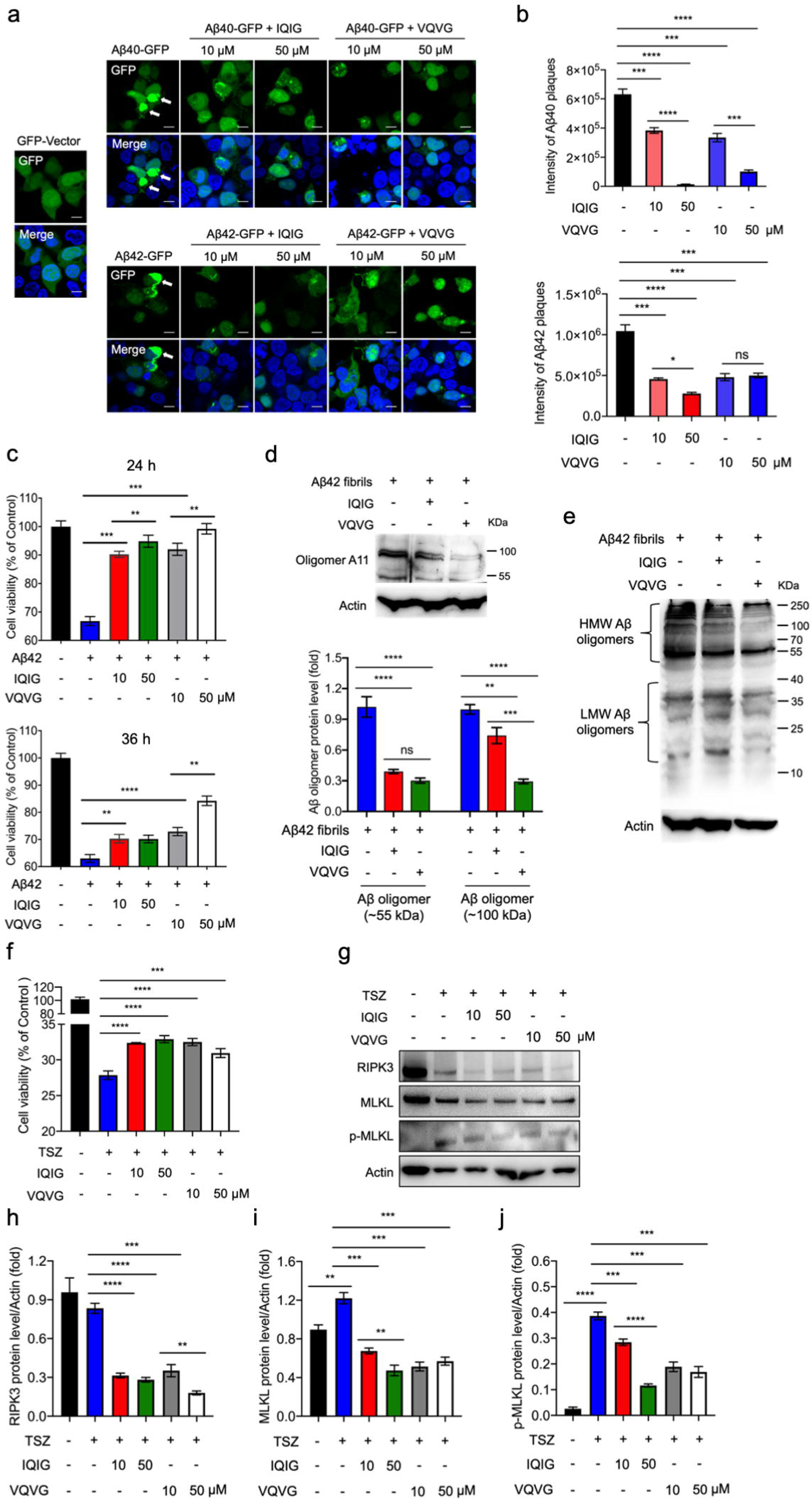
IQIG and VQVG inhibited neural cell death in response to Aβ aggregates. **a**. SH-SY5Y cells were transfected with GFP-tagged Aβ40 or Aβ42. Transfected cells were treated with or without different concentrations of IQIG and VQVG peptides for 24 h, followed by confocal microscopy analysis. Cells were fixed and stained with DAPI. The white arrows pointed to Aβ puncta. Scale bars, 10 μm. **b**. The relative intensity of Aβ40 and Aβ42 plaques was quantified by densitometry. **c**. The peptides IQIG and VQVG decreased Aβ aggregates-induced neural cell death. Human neuroblastoma SH-SY5Y cells were treated with various concentrations of IQIG or VQVG for 24-36 h. Aggregated Aβ42 was used as the positive control for inducing neural cell death. Cell viability was checked by the CCK8 assay and was normalized to that of the DMSO-treated cells. **d**. Cellular expression levels of indicated proteins in SH-SY5Y cells stimulated with Aβ42 aggregates in the presence or the absence of 10 μM IQIG or VQVG for 24 h. Western blot analysis was performed with an antibody specific for Aβ oligomer (anti-amyloidogenic protein oligomer A11). Quantification of the results for ∼55 kDa and ∼100 kDa Aβ oligomers were shown in the bottom panels. **e**. Western blot analysis was performed with an antibody for total Aβ protein (anti-Aβ) in SH-SY5Y cells treated as described in **d**. Aβ oligomerization was presented as high molecular weight oligomers (> 55 kDa) and low molecular weight oligomers (< 55 kDa). **f**. L929 cells were stimulated with 10 ng/ml TNFα, 100 nM Smac mimetic, and 10 μM Z-VAD (TSZ) in the presence or the absence of IQIG and VQVG for 4 h and subjected to the CCK8 assay for detecting cell viability (left). **g-j**. Western blot results for RIPK3, MLKL, and p-MLKL in L929 cells treated as described in **f**. Quantifications of the results for RIPK3, MLKL, and p-MLKL in treated or untreated cells were shown in **h-j**. The error bars represented the standard deviations (SD). One-way analysis of variance was performed (* *p* < 0.05, ** *p* < 0.01 and *** *p* < 0.001). All data are representative of three independent experiments.

Next, levels of Aβ oligomers in SH-SY5Y neuroblastoma cells were detected by the oligomer-specific Aβ antibody, A11, and then subjected to a semi-quantification analysis. It turned out that A11-immunoreactive oligomers (∼55 kDa and ∼100 kDa) were significantly reduced with IQIG or VQVG treatments (Fig. 3d). Furthermore, western blot analysis with the anti-Aβ antibody demonstrated that VQVG and IQIG treatments dramatically decreased both high molecular weight (HMW) and low molecular weight (LMW) Aβ oligomers (Fig. 3e). All of these data are in good agreement with the ThT fluorescence and TEM results, indicating that IQIG and VQVG could disassembly Aβ oligomers and fibrils, therefore reducing Aβ aggregates induced cell death.

Furthermore, we found that IQIG and VQVG could directly inhibit the polymerization of RIPK1 *in vitro* (Fig. S2a-S2b). To test the protective effects of IQIG and VQVG against necroptosis, we treated mouse fibroblast L929 cells with TSZ (TNFα/Smac/ZVAD) in the presence or absence of the tetrapeptides and then detected cell viability by the CCK8 assay. Cell viability significantly decreased in all cultured cells treated with IQIG or VQVG (Fig. 3f). In addition, the necroptotic markers RIPK3 and p-MLKL increased in the presence of TSZ, but decreased with IQIG or VQVG treatments, indicating that the tetrapeptides functioned in blocking TNFα-induced necroptosis (Fig. 3g-3i). Therefore, IQIG and VQVG could prevent TNFα-induced necroptosis and reduce Aβ aggregates triggered cell death.

### IQIG and VQVG increase learning and memory in aged APP/PS1 mice

The inhibitory effects of IQIG and VQVG on Aβ-induced cytotoxicity suggested a potential application for treating Aβ-related learning and memory deficits. Therefore, we examined IQIG and VQVG on spatial working memory and hippocampal memory using the classical aged APP^swe^/PS1^ΔE^^9^ (APP/PS1) double-transgenic mouse model ^39^. The APP/PS1 mice produced elevated levels of human Aβ because of the expression of mutant human APP and PS1, causing an AD-like phenotype starting at five months of age while cognitive impairment becomes observable around 6-8 months of age ^39^. After a dose-optimization trial of peptides, we intracerebroventricularly administered IQIG (10 mg/kg/mouse, n=17), VQVG (20 mg/kg/mouse, n=17), or vehicle (n=13) only (1 time/week) for three weeks to 6-month-old APP/PS1 mice until the age of 7 months (Fig. 4a). The 6-month-old wild type mice treated by vehicle (n=8) was administered at the same way and set as control.

**Figure 4.**
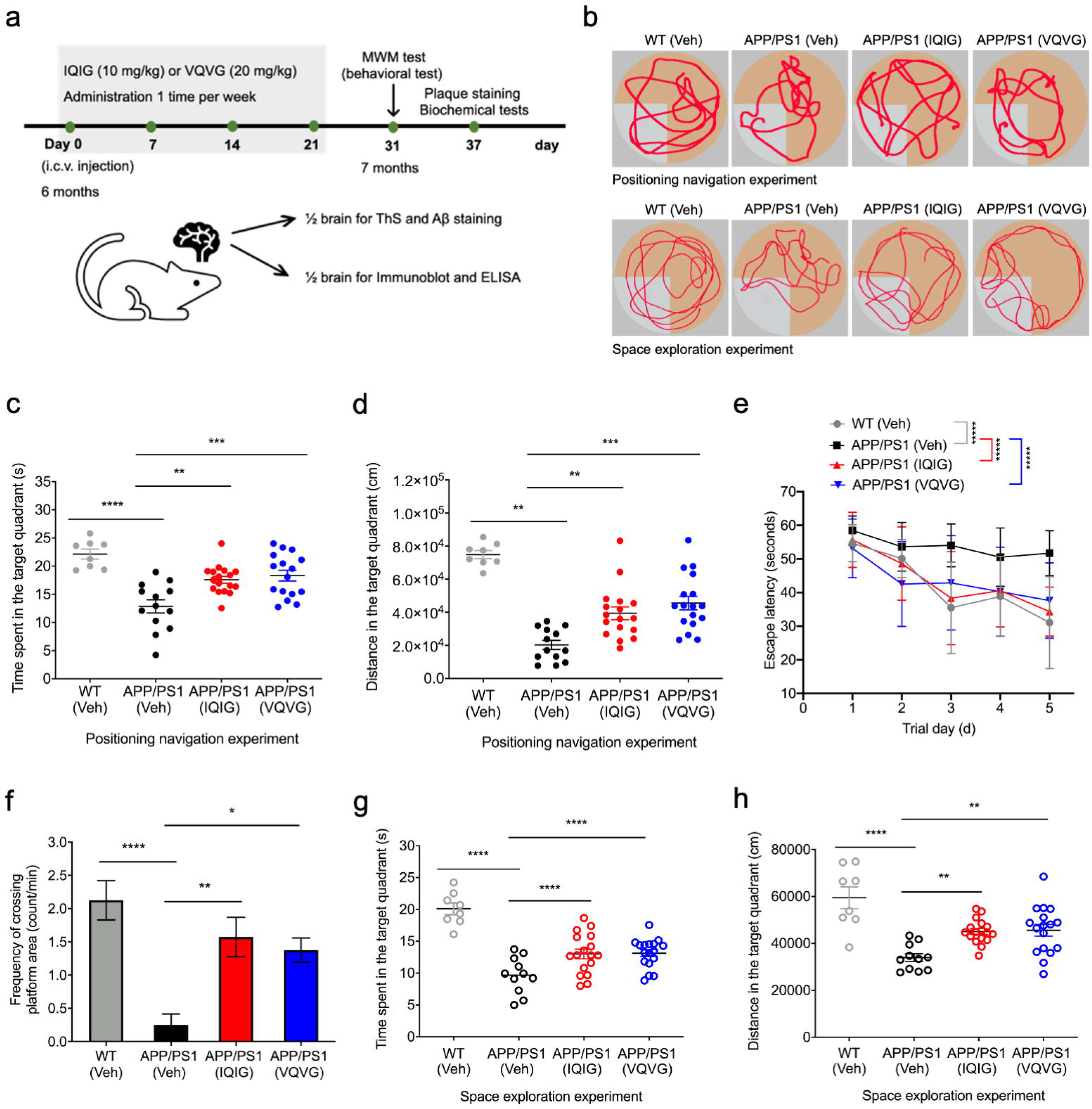
IQIG and VQVG improved the learning and memory abilities of the transgenic APP/PS1 mice. **a**. Schedule of IQIG and VQVG administration. IQIG (10 mg/kg, *n*=17), VQVG (20 mg/kg, *n*=17), or vehicle (PBS, *n*=13) was injected into 6-month-old male APP/PS1 mice in the intracerebroventricular region for three weeks (one time per week). The wild type mice treated with vehicle (PBS, *n*=8) was administrated at the same way. Acquisition tests were performed four times per day for seven days. The probe test was conducted on the eighth day without the platform. **b.** Representative path trackings in the MWM test with the hidden platform. **c**. Time spent in the platform quadrant in the probe test. **d**. Swim distance in the probe test. **e**. Hidden platform test in the MWM test. **f**. The target zone frequency in the probe test. **g**. Time spent in the platform quadrant in the space exploration experiment. **h**. Swim distance in the space exploration experiment. The error bars represented the standard deviations (SD). One-way or two-way analysis of variance was performed (* *p* < 0.05, ** *p* < 0.01 and *** *p* < 0.001). All data are representative of three independent experiments.

There were no obvious side effects (e.g., death of mice or significant reduction in body weight compared to control) observed during the experiment (Fig. S3e). Spatial learning and positioning navigation were performed using the Morris water maze (MWM) method, in which mice were placed in a circular water pool and tasked with finding a submerged, invisible escape platform (Fig. 4a-4b). The positioning navigation experiment was used to record and compare the movement distance and time of the mice to the platform in continuous training for five days; the space exploration experiment was used to record the time for the mice to reach the original platform and the times of crossing the original platform in the test.

APP/PS1 mice treated with the IQIG or VQVG peptide significantly increased the learning and navigation capabilities of untreated APP/PS1 mice (Fig. 4c-4d). In addition, escape latencies of mice treated with IQIG significantly decreased on days 3 (38.31 ± 4.36 s, *P* = 0.03) and 5 (36.05 ± 2.12 s, *P* = 0.0008), while those of mice treated with VQVG decreased markedly on day 5 (36.52 ± 3.39 s, *P* = 0.0016) (Fig. 4e). Mice treated with peptides also had higher frequencies of crossing platform area than those of the control APP/PS1 mice group (Fig. 4f), evidenced by 1.57 ± 0.30 counts/min (*P* = 0.0179) for the IQIG group, 1.38 ± 0.18 (*P* = 0.0472) for the VQVG group, and 0.25 ± 0.16 for the control group. Also, peptide-treated mice spent significantly more time and had a higher percentage of swimming paths in the target quadrant than in the control APP/PS1 mice group (Fig. 4g-4h). Overall, the data indicated that peptides IQIG and VQVG could improve cognition and alleviate AD symptoms in APP/PS1 mice.

### IQIG and VQVG reduce Aβ plaques in the cortex and hippocampus of aged APP/PS1 mice

To further identify whether IQIG and VQVG alleviate the cognitive impairments by directly reducing Aβ plaques, the amyloid-specific dye thioflavin S (ThS) and the anti-Aβ antibody were used to visualize the cryosections of brain tissues. We found that ThS-positive plaques were significantly reduced in the brains of intracerebroventricularly IQIG- or VQVG-injected mice (Fig. 5a-5b). Also, Aβ-positive plaques were spread throughout the entire cortex in the APP/PS mice. At the same time, a small amount of Aβ could be detected in the brains of IQIG-injected mice (Fig. 5a, 5c-5e). Quantification analysis showed that Aβ plaques dramatically decreased in the hippocampus and cortex regions of the IQIG- or VQVG-injected mice (Fig. 5d-5e). As hypophosphorylated tau (p-tau) is a histopathological hallmark of AD ^2^, we also detected the p-tau expression level in the APP/PS1 mice. The results showed that IQIG and VQVG treatments significantly reduced the abundance of p-tau tangles in the hippocampus (Fig. 5f and 5h).

**Figure 5.**
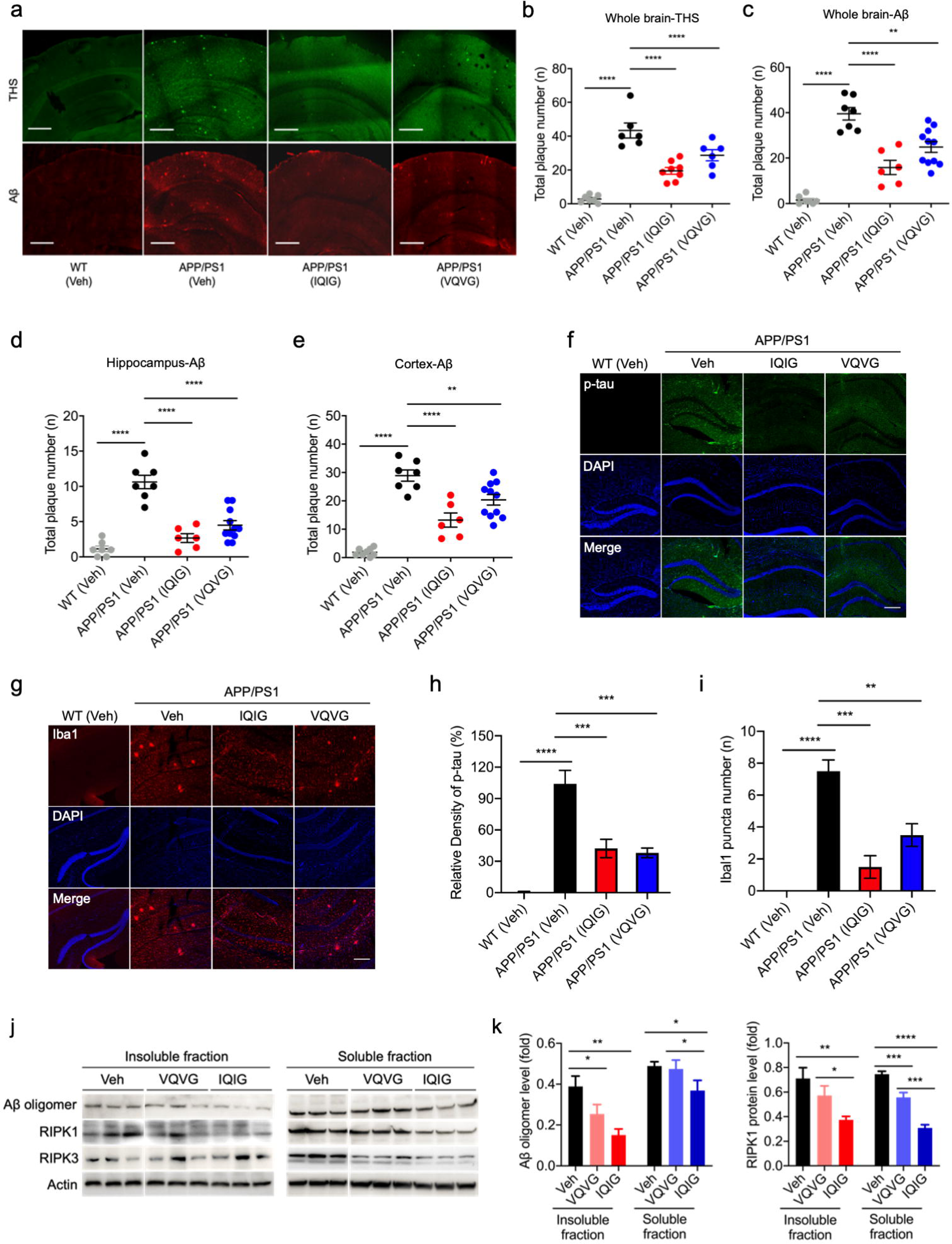
IQIG and VQVG reduced Aβ plaques, p-tau tangles, and microglial plaques in the brains of the APP/PS1 mice. The APP/PS1 or wild type mice subjected to behavioral tests were used for brain analyses. **a**. The hippocampal and cortex regions were stained with the amyloid-specific binding dye ThS or the anti-Aβ antibody (scale bars, 500 μm). **b**. IQIG and VQVG decreased ThS-positive plaques the APP/PS1 mice. **c-e**. Amyloid plaque numbers were quantified in the total brain (**c**), hippocampus (**d)**, and cortex regions (**e**). **f-i**. Immunostaining for p-tau (**f**), Iba1 (**g**), and DAPI after administration of vehicle, IQIG, or VQVG in the hippocampus of the APP/PS1 mice, or wild type mice (*n* = 6 for each group). Scale bars, 200 μm. Quantification results of p-tau and Iba1 are shown in **h** and **i**. **j-k**. Western blotting analyses of Aβ oligomer, RIPK1, and RIPK3 in the insoluble hippocampal (**left**) and soluble hippocampal (**right**) lysates. The quantification results of Aβ oligomer expression levels are shown in **k**. Data are denoted as mean ± SD, One-way analysis of variance was performed (* *p* < 0.05, ** *p* < 0.01, *** *p* < 0.001 and **** *p* < 0.0001). All data are representative of three independent experiments.

A number of ionized calcium-binding adaptor 1 (Iba1) specks were detected in six months old APP/PS1 mice, exhibiting morphological signs of microglial activation as previous reports ^40^ (Fig. S3a). After IQIG and VQVG treatments, the microglia puncta decreased dramatically (Fig. 5g and 5i). Previous reports showed that soluble Aβ oligomer is particularly neurotoxic and indicates central brain dysfunction in AD^41^. Inhibition of RIPK1 kinase activity can reduce neuroinflammation mediated by microglia ^40^. Therefore, we checked the protein expression levels of Aβ, RIPK1, and RIPK3. The results showed that RIPK1 and Aβ that detected by the specific anti-Aβ oligomer A11 antibody, dramatically decreased in the soluble and insoluble hippocampal lysates of IQIG-injected mice (Fig. 5j-5k). Thus, the peptides might have multiple mechanisms on AD pathology, including directly targeting and regulating RIPKs and Aβ/tau aggregations.

Besides, compared to the three months old mice, the APP/PS1 mice at six months revealed a significant increase in RIPK1 and RIPK3 expression (Fig. S3b); however, both RIPK1 and RIPK3 did not co-localize with Aβ, similar to the previous reports (Fig. S3b) ^40^. Also, the anti-inflammatory cytokines’ expression levels, including IL-4, IL-10, and IL-13, dramatically increased in the IQIG- or VQVG-injected mice (Fig. S3c-S3d). These results showed that treatments with IQIG or VQVG could alleviate Aβ aggregates-induced neuroinflammation. Taken together, IQIG and VQVG could decrease the oligomerization of Aβ, alleviate Aβ plaques, and inhibit neuroinflammation in APP/PS1 mice.

### IQIG and VQVG restored cellular homeostasis at the single-cell level in the hippocampus of aged APP/PS1 mice

To explore how IQIG and VQVG peptides alleviate AD symptoms and improve cognition in aged APP/PS1 mice from single cell level, we extracted hippocampus region of IQIG-injected, VQVG-injected, and vehicle-injected (Ctrl) APP/PS1 mice and performed single-cell RNA sequencing (scRNA-seq) based on 10X Chromium platform (Fig. 6a). Based on 8,934 cells from 9 mice (3 mice for each group), we identified 30 cell subtypes in total, which contain 11 glutamatergic neuron subtypes, 6 GABAergic neuron subtypes, 1 Cajal-Retzius cell (CR) subtype, 2 microglia subtypes, 2 astrocyte subtypes, 4 oligodendrocyte subtypes, 2 oligodendrocyte progenitor cells (OPCs) subtypes, 1 vascular/leptomeningeal cell (VLMC) subtypes (Fig. 6b and S4). The two microglia subtypes can be further identified as homeostatic microglia (Micro_0) and disease-associated microglia (DAM, Micro_1) based on their transcriptomic profiles (Fig. 6c-6d). Our results demonstrate a significant increase in cell proportions of GABA_3, GABA_5 GABAergic neurons, and Oligo_1 oligodendrocytes, while a noteworthy decrease was observed for GLU_0, GLU_1 glutamatergic neurons, Oligo_3 oligodendrocytes, and Astro_0 astrocytes, in both IQIG- and VQVG-treated mice when compared to the control group (Fig. 6e, Table S1). The neuronal loss of GABAergic neurons and disrupted neural network activity were reported in the hippocampus of both human AD patients and AD mouse models as a hallmark of AD ^42–46^. The loss of GABAergic neurons can result in deficits within the inhibitory network, leading to hyperexcitability and subsequent dysregulation of neural circuits. Such disturbances can further culminate in cognitive decline ^47^. The orchestration between inhibitory and excitatory neurons is pivotal for maintaining neural homeostasis and function. The increase of GABAergic neuron subtypes and decrease of glutamatergic neuron subtypes in the treated groups indicates the restored homeostasis of the neuronal network in IQIG- and VQVG-treated mice, which further supports our experimental observation of improved cognitive function in treated mice.

**Figure 6.**
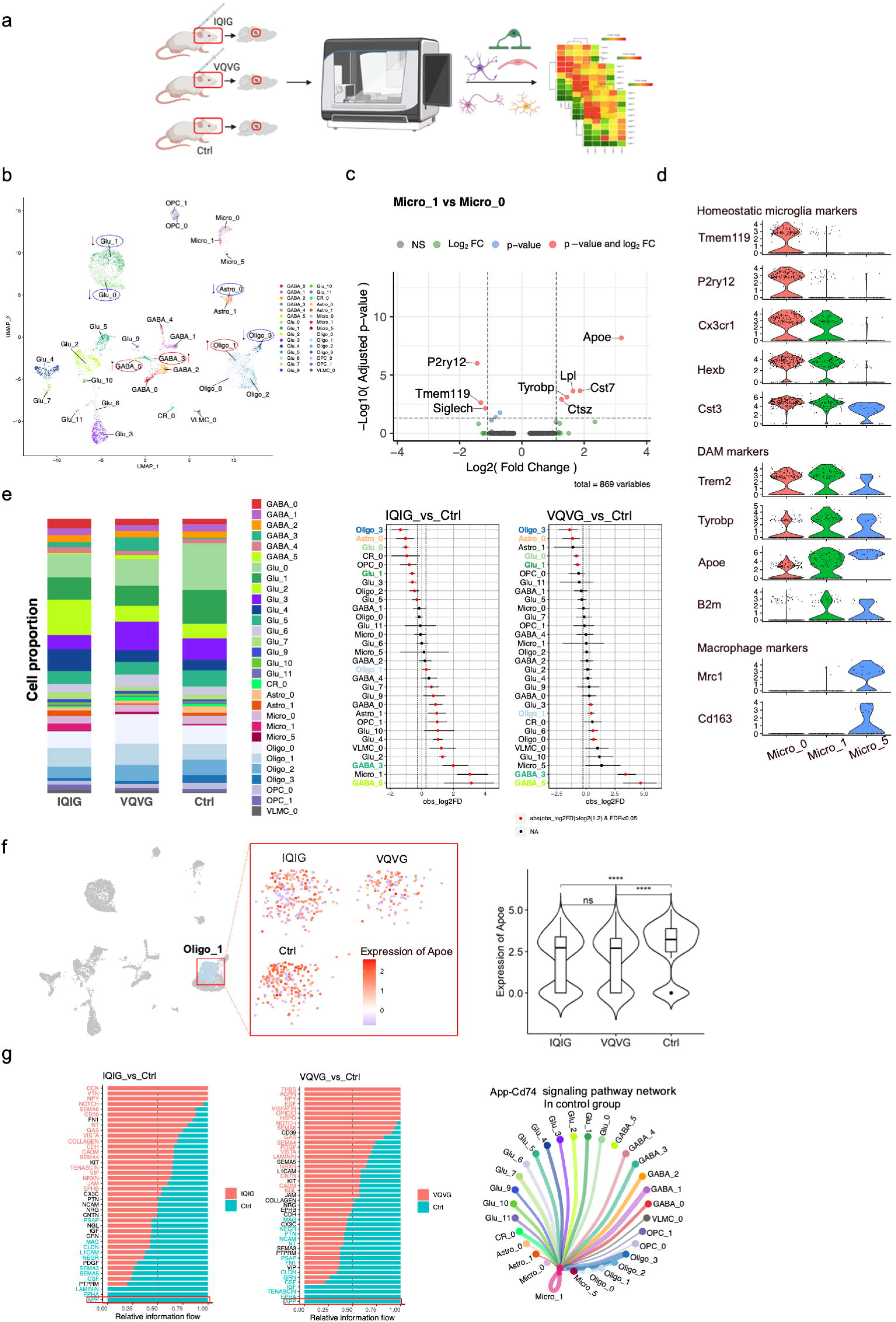
Comparative analysis of single-cell transcriptomic profiles between the peptide treatment and control groups of aged APP/PS1 mice. **a**. Schematic representation of the experimental design for single-cell RNA sequencing. **b**. Uniform Manifold Approximation and Projection (UMAP) plot illustrating 30 cell subtypes. Colors represent different subtypes. Prefixes for subtype names are as follows: Glu=Glutamatergic, GABA=GABAergic, CR=Cajal-Retzius cell, Oligo=Oligodendrocyte, VLMC=Vascular/Leptomeningeal cell, Astro=Astrocyte, Micro=Microglia, OPC=Oligodendrocyte progenitor cell. Subtypes encircled in red, or blue indicate consistent increased or decreased proportional changes, respectively, in both IQIV_vs_Ctrl and VQVG_vs_Ctrl contrasts. **c-d**. Evidence of microglia subtype annotation. **c**. Volcano plot showing the differential expressed genes between Micro_0 and Micro_1. **d**. Expression of homeostatic microglia, disease-associated microglia (DAM), and macrophage markers in Micro_0, Micro_1, and Micro_5. **e**. Bar and dot plots illustrate proportion changes of each cell type post-treatment. In the dot plot, a red dot denotes a significant proportion change, while colored and bolded y-axis labels highlight cell types with consistent proportional shifts in both IQIV_vs_Ctrl and VQVG_vs_Ctrl contrasts. **f**. Dot plot and violin plot showing the expression of the Apoe gene in each group. **g**. Bar plot showing the relative information flow of each signal pathway. The APP signal pathway can only be detected in the control group and is highlighted by red rectangles. Network plot showing the APP signal pathway strength between cell types in the control group.

The expression level of the known AD risk gene, *Apoe*, decreased significantly in the OLIGO_1 oligodendrocyte subtype in both IQIG- and VQVG-treated mice compared to the control mice (Fig. 6f and Table S2). *Apoe* gene encodes apolipoprotein E, which is associated with cholesterol and lipid metabolism ^48–50^. Accumulated evidence in recent years of research has emphasized the complex function of the *APOE* gene and the interaction between the apoE protein and Aβ in the pathology of AD ^51–53^. The isoform APOE4 can impair myelination in oligodendrocytes through cholesterol dysregulation based on a previous study ^54^. Our scRNA-seq results show the decreased Apoe expression in the Oligo_1 subtype of the treated groups, which suggests that reducing Aβ may decrease the Apoe level in specific oligodendrocyte subtype, which may increase axonal myelination and improve learning and memory in mice (Fig. 6).

Through cell-cell communication analysis by CellChat, we observed increased total cell-cell interaction strength in the IQIG-treated, VQVG-treated mice compared with the control APP/PS1 mice (Fig. S6a). Interestingly, the APP relative information flow showed specificity between DAM (disease-associated microglia) and other cell types in the control group, which indicated the IQIG and VQVG treatment could efficiently decrease the APP-related signaling pathway (Fig. 6g and S6b, Table S3). Even though there is no significant change in the cell proportion of microglia (Fig. S5), we observed a consistent changing pattern of cell-cell interaction in both contrasts (i.e., IQIG-treated vs. control groups, and VQVG-treated vs. control groups) (Fig. S7 and Table S4). The consistent changing pattern shows decreased interaction strength between DAM and other cells (for example, oligodendrocytes, astrocytes, OPC_1, Glu_7, Glu_10, Glu_11), while increased interaction between the homeostatic microglia and specific cell subtypes, such as GABA_5, GABA_4 and OPC_0, in both the IQIG- and VQVG-treated group compared with the control group (Fig. S8-S10 and Table S5-S6). The changing pattern of cell-cell interaction supports our previous claim that IQIG and VQVG peptide treatment decreased microglia-mediated neuroinflammation.

## DISCUSSION

Alzheimer’s disease (AD) is characterized by a specific neuropathological change pattern, including extracellular Aβ deposits, intracellular neurofibrillary tangles, granulovacuolar degeneration representing cytoplasmic vacuolar lesions, synapse dysfunction, and neuronal loss ^55^. The connection between necroptosis and neuronal damage in AD has been suggested by demonstrating a protective effect of the necroptosis inhibitor on brain injury ^56^. The deficiency of RIPK3 alleviates the loss of hippocampal neurons after intracerebroventricular injection of TNF-α ^57^. Here, we discovered that the RIPK1/RIPK3 core RHIM peptides inhibited and disaggregated Aβ fibrils and plaques *in vitro* and *in vivo*. Also, IQIG and VQVG treatments blocked Aβ-induced neuronal cell death (Fig. 1-3). Furthermore, the mice treated with IQIG or VQVG had a decreased escape latency in the acquisition phase and increased swimming times in the target quadrant, indicating that the peptides could improve long-time spatial learning and memory capacities in the APP/PS1 mice (Fig. 4). Moreover, the intracerebroventricular administration of IQIG or VQVG exhibited a histological reduction of the amyloid burden, as well as p-tau tangles and Iba1 plaques in the APP/PS1 mice (Fig. 5).

Nec-1 can reduce the extent of injury or block microglial dysfunction in various animal models ^58, 59^, but can not directly inhibit neuronal cell death induced by Aβ, suggesting that Nec-1 does not directly target Aβ or directly inhibit neuronal cell death ^40, 60, 61^. Our results showed that IQIG and VQVG could directly bind to Aβ40 or Aβ42 and efficiently inhibited Aβ aggregation (Fig. 1 and S1); however, the protective effects for the two peptides were not on the same level. IQIG had a better capacity for decreasing Aβ plaques and p-tau tangles and alleviating cognitive impairments in the APP/PS1 mice (Fig. 5). As the pleiotropic RIPK1 acts in multiple signaling pathways, we speculated that the RIPK1-derived peptide IQIG might function more efficiently in inhibiting both TNFα-induced cell necrosis and Aβ aggregates in AD.

Mice deficient in NLRP3 inflammasome (*Nlrp3^-/-^*), required for the caspase-1-dependent secretion of IL-1β and IL-18, have reduced inflammation and improved cognition and memory ^62–64^. Also, the increased level of TNF-α is detected in the brain and plasma in AD patients and mouse models of AD ^65, 66^. Moreover, multiple inflammatory cytokines contribute to the activation of microglia and Aβ generation and deposition ^67–69^. Both IQIG and VQVG treatments increased the expression level of anti-inflammatory cytokines (IL4, IL-10, and IL-13) in the hippocampus, indicating a positive correlation between the anti-inflammatory cytokine production and Aβ clearance (Fig. S3) ^56^.

Our scRNA-seq analysis results clearly demonstrate that treatment with IQIG and VQVG peptides restored the cellular homeostasis in the hippocampus of aged APP/PS1 mice, which leads to a significant increase in two GABAergic neuron subtypes, while simultaneously decreasing two glutamatergic neuron subtypes in the hippocampus of aged APP/PS1 mice (Fig. 6a-6e). Additionally, we observed a decrease in *Apoe* expression level and reduced APP-related and DAM-related cell-cell communication (Fig. 6f-6g). These results provide strong evidence at the single-cell level, supporting our previous claim that IQIG and VQVG peptides effectively alleviate pathological changes and enhance cognitive and memory function in AD mice.

In summary, the RIPK1/RIPK3 core RHIM peptides could directly bind to Aβ, inhibit its polymerization, and disassemble Aβ preformed fibrils. Besides, we proved the protective effects of IQIG and VQVG in the APP/PS1 transgenic mice, which might provide a new strategy for preventing and treating AD. As the activation of necroptosis is also reported in other neurodegenerative diseases, including PD, ALS, and multiple sclerosis ^12, 17, 18, 70^, it remains open whether the peptides could inhibit neuronal loss and dysfunction in these diseases. These exciting possibilities should be examined in future investigations.

## CONSENT STATEMENT

The study did not involve with human subjects and the consent was unnecessary.

## ACKNOWLEDGMENTS

We thank Dr. Y Mei and Dr. B Lu (Fudan University) for helping with the mouse behaviour tests, and the AMSP (aged mice sharing project) (SIBCB, CAS) for providing aged mice.

## AUTHOR CONTRIBUTIONS

J.L. and H.W. conceived and designed the study; W.G., B.G., S.W., D.W., P.M., L.G., C.L., Y.L., X.L., J.P., J.H., Y.D., X.C., H.S., H.Y., S.L., Q.Z., B.X., and F.H. performed the experiments and analyzed the data. C.M. analyzed the scRNA-seq data. J.L. and W.G. analyzed the data and wrote the manuscript. All authors discussed the results and commented on the manuscript.

## DATA AND MATERIALS AVAILABILITY

All single-cell RNA sequencing data are available from the Gene Expression Omnibus (GEO) under the accession number GSE242308. All data needed to evaluate the conclusions in the paper are present in the paper and/or the Supplementary Materials. Additional data related to this paper may be requested from the corresponding author.

## ETHICS DECLARATIONS

All animal experiments were approved in accordance with IACUC regulations by the Scientific Investigation Committee of the School of Life Sciences, Fudan University (2019-JS-011).

## REFERENCES

1 Lane, C. A., Hardy, J. & Schott, J. M. Alzheimer’s disease. Eur J Neurol 25, 59–70, doi:10.1111/ene.13439 (2018).

2 Long, J. M. & Holtzman, D. M. Alzheimer Disease: An Update on Pathobiology and Treatment Strategies. Cell 179, 312–339, doi:10.1016/j.cell.2019.09.001 (2019).

3 Tolar, M., Abushakra, S. & Sabbagh, M. The path forward in Alzheimer’s disease therapeutics: Reevaluating the amyloid cascade hypothesis. Alzheimers Dement, doi:10.1016/j.jalz.2019.09.075 (2020).

4 Lee, S. J., Nam, E., Lee, H. J., Savelieff, M. G. & Lim, M. H. Towards an understanding of amyloid-beta oligomers: characterization, toxicity mechanisms, and inhibitors. Chem Soc Rev 46, 310–323, doi:10.1039/c6cs00731g (2017).

5 Kim, H. Y. et al. EPPS rescues hippocampus-dependent cognitive deficits in APP/PS1 mice by disaggregation of amyloid-beta oligomers and plaques. Nat Commun 6, 8997, doi:10.1038/ncomms9997 (2015).

6 Hefti, F. et al. The case for soluble Abeta oligomers as a drug target in Alzheimer’s disease. Trends Pharmacol Sci 34, 261–266, doi:10.1016/j.tips.2013.03.002 (2013).

7 Koss, D. J. et al. Soluble pre-fibrillar tau and beta-amyloid species emerge in early human Alzheimer’s disease and track disease progression and cognitive decline. Acta Neuropathol 132, 875–895, doi:10.1007/s00401-016-1632-3 (2016).

8 Chen, Z. L., Singh, P. K., Calvano, M., Norris, E. H. & Strickland, S. A possible mechanism for the enhanced toxicity of beta-amyloid protofibrils in Alzheimer’s disease. Proc Natl Acad Sci U S A 120, e2309389120, doi:10.1073/pnas.2309389120 (2023).

9 Harris, E. Alzheimer Drug Lecanemab Gains Traditional FDA Approval. JAMA 330, 495, doi:10.1001/jama.2023.12548 (2023).

10 Seidler, P. M. et al. Structure-based inhibitors of tau aggregation. Nat Chem 10, 170–176, doi:10.1038/nchem.2889 (2018).

11 Neddenriep, B. et al. Short Peptides as Inhibitors of Amyloid Aggregation. Open Biotechnol J 5, 39–46, doi:10.2174/1874070701105010039 (2011).

12 Caccamo, A. et al. Necroptosis activation in Alzheimer’s disease. Nat Neurosci 20, 1236–1246, doi:10.1038/nn.4608 (2017).

13 Degterev, A., Ofengeim, D. & Yuan, J. Targeting RIPK1 for the treatment of human diseases. Proc Natl Acad Sci U S A 116, 9714–9722, doi:10.1073/pnas.1901179116 (2019).

14 Koper, M. J. et al. Necrosome complex detected in granulovacuolar degeneration is associated with neuronal loss in Alzheimer’s disease. Acta Neuropathol 139, 463–484, doi:10.1007/s00401-019-02103-y (2020).

15 Cho, Y. S. et al. Phosphorylation-driven assembly of the RIP1-RIP3 complex regulates programmed necrosis and virus-induced inflammation. Cell 137, 1112–1123, doi:10.1016/j.cell.2009.05.037 (2009).

16 He, S. et al. Receptor interacting protein kinase-3 determines cellular necrotic response to TNF-alpha. Cell 137, 1100–1111, doi:10.1016/j.cell.2009.05.021 (2009).

17 Onate, M. et al. The necroptosis machinery mediates axonal degeneration in a model of Parkinson disease. Cell Death Differ 27, 1169–1185, doi:10.1038/s41418-019-0408-4 (2020).

18 Ito, Y. et al. RIPK1 mediates axonal degeneration by promoting inflammation and necroptosis in ALS. Science 353, 603–608, doi:10.1126/science.aaf6803 (2016).

19 Ofengeim, D. et al. Activation of necroptosis in multiple sclerosis. Cell Rep 10, 1836–1849, doi:10.1016/j.celrep.2015.02.051 (2015).

20 Lloyd, A. F. et al. Central nervous system regeneration is driven by microglia necroptosis and repopulation. Nat Neurosci 22, 1046–1052, doi:10.1038/s41593-019-0418-z (2019).

21 Grootjans, S., Vanden Berghe, T. & Vandenabeele, P. Initiation and execution mechanisms of necroptosis: an overview. Cell Death Differ 24, 1184–1195, doi:10.1038/cdd.2017.65 (2017).

22 Chen, J., Kos, R., Garssen, J. & Redegeld, F. Molecular Insights into the Mechanism of Necroptosis: The Necrosome As a Potential Therapeutic Target. Cells 8, doi:10.3390/cells8121486 (2019).

23 Li, J. et al. The RIP1/RIP3 necrosome forms a functional amyloid signaling complex required for programmed necrosis. Cell 150, 339–350, doi:10.1016/j.cell.2012.06.019 (2012).

24 Mompean, M. et al. The Structure of the Necrosome RIPK1-RIPK3 Core, a Human Hetero-Amyloid Signaling Complex. Cell 173, 1244–1253 e1210, doi:10.1016/j.cell.2018.03.032 (2018).

25 Zhang, Q. et al. Unnatural Peptide Assemblies Rapidly Deplete Cholesterol and Potently Inhibit Cancer Cells. J Am Chem Soc 146, 12901–12906, doi:10.1021/jacs.4c03101 (2024).

26 Hao, Y. et al. Integrated analysis of multimodal single-cell data. Cell 184, 3573–3587 e3529, doi:10.1016/j.cell.2021.04.048 (2021).

27 Yao, Z. et al. A taxonomy of transcriptomic cell types across the isocortex and hippocampal formation. Cell 184, 3222–3241 e3226, doi:10.1016/j.cell.2021.04.021 (2021).

28 Miller, S. A. et al. LSD1 and Aberrant DNA Methylation Mediate Persistence of Enteroendocrine Progenitors That Support BRAF-Mutant Colorectal Cancer. Cancer Res 81, 3791–3805, doi:10.1158/0008-5472.CAN-20-3562 (2021).

29 Paravastu, A. K., Leapman, R. D., Yau, W. M. & Tycko, R. Molecular structural basis for polymorphism in Alzheimer’s beta-amyloid fibrils. Proc Natl Acad Sci U S A 105, 18349–18354, doi:10.1073/pnas.0806270105 (2008).

30 Petkova, A. T., Yau, W. M. & Tycko, R. Experimental constraints on quaternary structure in Alzheimer’s beta-amyloid fibrils. Biochemistry 45, 498–512, doi:10.1021/bi051952q (2006).

31 Fandrich, M., Schmidt, M. & Grigorieff, N. Recent progress in understanding Alzheimer’s beta-amyloid structures. Trends Biochem Sci 36, 338–345, doi:10.1016/j.tibs.2011.02.002 (2011).

32 Ghosh, U., Thurber, K. R., Yau, W. M. & Tycko, R. Molecular structure of a prevalent amyloid-beta fibril polymorph from Alzheimer’s disease brain tissue. Proc Natl Acad Sci U S A 118, doi:10.1073/pnas.2023089118 (2021).

33 Nelson, R. & Eisenberg, D. Recent atomic models of amyloid fibril structure. Curr Opin Struct Biol 16, 260–265, doi:10.1016/j.sbi.2006.03.007 (2006).

34 Armiento, V., Spanopoulou, A. & Kapurniotu, A. Peptide-Based Molecular Strategies To Interfere with Protein Misfolding, Aggregation, and Cell Degeneration. Angew Chem Int Ed Engl 59, 3372–3384, doi:10.1002/anie.201906908 (2020).

35 Blanchard, B. J. et al. Efficient reversal of Alzheimer’s disease fibril formation and elimination of neurotoxicity by a small molecule. Proc Natl Acad Sci U S A 101, 14326–14332, doi:10.1073/pnas.0405941101 (2004).

36 He, C., Hou, Y., Han, Y. & Wang, Y. Disassembly of amyloid fibrils by premicellar and micellar aggregates of a tetrameric cationic surfactant in aqueous solution. Langmuir 27, 4551–4556, doi:10.1021/la200350j (2011).

37 Li, J., Zhu, M., Manning-Bog, A. B., Di Monte, D. A. & Fink, A. L. Dopamine and L-dopa disaggregate amyloid fibrils: implications for Parkinson’s and Alzheimer’s disease. FASEB J 18, 962–964, doi:10.1096/fj.03-0770fje (2004).

38 Du, W. J. et al. Brazilin inhibits amyloid beta-protein fibrillogenesis, remodels amyloid fibrils and reduces amyloid cytotoxicity. Sci Rep 5, 7992, doi:10.1038/srep07992 (2015).

39 Heneka, M. T. et al. NLRP3 is activated in Alzheimer’s disease and contributes to pathology in APP/PS1 mice. Nature 493, 674–678, doi:10.1038/nature11729 (2013).

40 Ofengeim, D. et al. RIPK1 mediates a disease-associated microglial response in Alzheimer’s disease. Proc Natl Acad Sci U S A 114, E8788–E8797, doi:10.1073/pnas.1714175114 (2017).

41 Benilova, I., Karran, E. & De Strooper, B. The toxic Abeta oligomer and Alzheimer’s disease: an emperor in need of clothes. Nat Neurosci 15, 349–357, doi:10.1038/nn.3028 (2012).

42 Jimenez-Balado, J. & Eich, T. S. GABAergic dysfunction, neural network hyperactivity and memory impairments in human aging and Alzheimer’s disease. Semin Cell Dev Biol 116, 146–159, doi:10.1016/j.semcdb.2021.01.005 (2021).

43 Levenga, J. et al. Tau pathology induces loss of GABAergic interneurons leading to altered synaptic plasticity and behavioral impairments. Acta Neuropathol Commun 1, 34, doi:10.1186/2051-5960-1-34 (2013).

44 Najm, R., Jones, E. A. & Huang, Y. Apolipoprotein E4, inhibitory network dysfunction, and Alzheimer’s disease. Mol Neurodegener 14, 24, doi:10.1186/s13024-019-0324-6 (2019).

45 Petrache, A. L. et al. Aberrant Excitatory-Inhibitory Synaptic Mechanisms in Entorhinal Cortex Microcircuits During the Pathogenesis of Alzheimer’s Disease. Cereb Cortex 29, 1834–1850, doi:10.1093/cercor/bhz016 (2019).

46 Ramos, B. et al. Early neuropathology of somatostatin/NPY GABAergic cells in the hippocampus of a PS1xAPP transgenic model of Alzheimer’s disease. Neurobiol Aging 27, 1658–1672, doi:10.1016/j.neurobiolaging.2005.09.022 (2006).

47 Xu, Y., Zhao, M., Han, Y. & Zhang, H. GABAergic Inhibitory Interneuron Deficits in Alzheimer’s Disease: Implications for Treatment. Front Neurosci 14, 660, doi:10.3389/fnins.2020.00660 (2020).

48 Arnon, R., Sehayek, E., Vogel, T. & Eisenberg, S. Effects of exogenous apo E-3 and of cholesterol-enriched meals on the cellular metabolism of human chylomicrons and their remnants. Biochim Biophys Acta 1085, 336–342, doi:10.1016/0005-2760(91)90138-8 (1991).

49 Krimbou, L. et al. Molecular interactions between apoE and ABCA1: impact on apoE lipidation. J Lipid Res 45, 839–848, doi:10.1194/jlr.M300418-JLR200 (2004).

50 Marcel, Y. L., Vezina, C. & Milne, R. W. Cholesteryl ester and apolipoprotein E transfer between human high density lipoproteins and chylomicrons. Biochim Biophys Acta 750, 411–417, doi:10.1016/0005-2760(83)90047-4 (1983).

51 Martens, Y. A. et al. ApoE Cascade Hypothesis in the pathogenesis of Alzheimer’s disease and related dementias. Neuron 110, 1304–1317, doi:10.1016/j.neuron.2022.03.004 (2022).

52 Namba, Y., Tomonaga, M., Kawasaki, H., Otomo, E. & Ikeda, K. Apolipoprotein E immunoreactivity in cerebral amyloid deposits and neurofibrillary tangles in Alzheimer’s disease and kuru plaque amyloid in Creutzfeldt-Jakob disease. Brain Res 541, 163–166, doi:10.1016/0006-8993(91)91092-f (1991).

53 Raulin, A. C. et al. ApoE in Alzheimer’s disease: pathophysiology and therapeutic strategies. Mol Neurodegener 17, 72, doi:10.1186/s13024-022-00574-4 (2022).

54 Blanchard, J. W. et al. APOE4 impairs myelination via cholesterol dysregulation in oligodendrocytes. Nature 611, 769–779, doi:10.1038/s41586-022-05439-w (2022).

55 Caselli, R. J., Beach, T. G., Yaari, R. & Reiman, E. M. Alzheimer’s disease a century later. J Clin Psychiatry 67, 1784–1800, doi:10.4088/jcp.v67n1118 (2006).

56 Kaczmarek, A., Vandenabeele, P. & Krysko, D. V. Necroptosis: the release of damage-associated molecular patterns and its physiological relevance. Immunity 38, 209–223, doi:10.1016/j.immuni.2013.02.003 (2013).

57 Vandenabeele, P., Declercq, W., Van Herreweghe, F. & Vanden Berghe, T. The role of the kinases RIP1 and RIP3 in TNF-induced necrosis. Sci Signal 3, re4, doi:10.1126/scisignal.3115re4 (2010).

58 Oerlemans, M. I. et al. Inhibition of RIP1-dependent necrosis prevents adverse cardiac remodeling after myocardial ischemia-reperfusion in vivo. Basic Res Cardiol 107, 270, doi:10.1007/s00395-012-0270-8 (2012).

59 Zhuang, C. & Chen, F. Small-Molecule Inhibitors of Necroptosis: Current Status and Perspectives. J Med Chem 63, 1490–1510, doi:10.1021/acs.jmedchem.9b01317 (2020).

60 Yang, S. H. et al. A small molecule Nec-1 directly induces amyloid clearance in the brains of aged APP/PS1 mice. Sci Rep 9, 4183, doi:10.1038/s41598-019-40205-5 (2019).

61 Yang, S. H. et al. Nec-1 alleviates cognitive impairment with reduction of Abeta and tau abnormalities in APP/PS1 mice. EMBO Mol Med 9, 61–77, doi:10.15252/emmm.201606566 (2017).

62 Ising, C. et al. NLRP3 inflammasome activation drives tau pathology. Nature 575, 669–673, doi:10.1038/s41586-019-1769-z (2019).

63 Mangan, M. S. J. et al. Targeting the NLRP3 inflammasome in inflammatory diseases. Nat Rev Drug Discov 17, 588–606, doi:10.1038/nrd.2018.97 (2018).

64 Olsen, I. & Singhrao, S. K. Inflammasome Involvement in Alzheimer’s Disease. J Alzheimers Dis 54, 45–53, doi:10.3233/JAD-160197 (2016).

65 Decourt, B., Lahiri, D. K. & Sabbagh, M. N. Targeting Tumor Necrosis Factor Alpha for Alzheimer’s Disease. Curr Alzheimer Res 14, 412–425, doi:10.2174/1567205013666160930110551 (2017).

66 Shamim, D. & Laskowski, M. Inhibition of Inflammation Mediated Through the Tumor Necrosis Factor alpha Biochemical Pathway Can Lead to Favorable Outcomes in Alzheimer Disease. J Cent Nerv Syst Dis 9, 1179573517722512, doi:10.1177/1179573517722512 (2017).

67 Calsolaro, V. & Edison, P. Neuroinflammation in Alzheimer’s disease: Current evidence and future directions. Alzheimers Dement 12, 719–732, doi:10.1016/j.jalz.2016.02.010 (2016).

68 Domingues, C., da Cruz, E. S. O. A. B. & Henriques, A. G. Impact of Cytokines and Chemokines on Alzheimer’s Disease Neuropathological Hallmarks. Curr Alzheimer Res 14, 870–882, doi:10.2174/1567205014666170317113606 (2017).

69 Minter, M. R., Taylor, J. M. & Crack, P. J. The contribution of neuroinflammation to amyloid toxicity in Alzheimer’s disease. J Neurochem 136, 457–474, doi:10.1111/jnc.13411 (2016).

70 Re, D. B. et al. Necroptosis drives motor neuron death in models of both sporadic and familial ALS. Neuron 81, 1001–1008, doi:10.1016/j.neuron.2014.01.011 (2014).

