## Supplementary Materials for "Peptides alleviate cognitive impairment by inhibiting and disassembling amyloid-β aggregates in Alzheimer’s disease"

**SUPPLEMENTARY FIGURES**

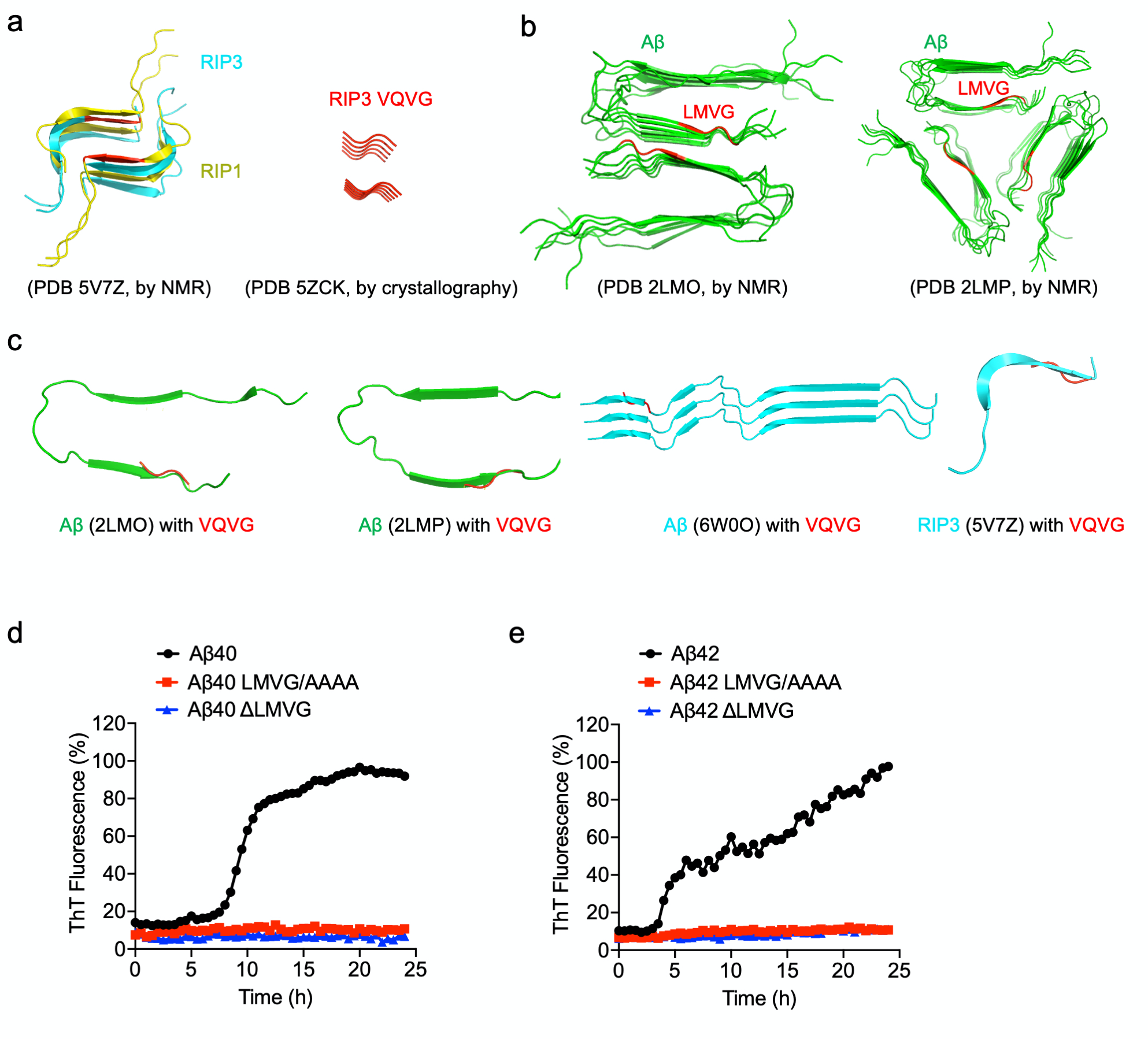

**Figure S1**. Protein structures and structural alignment of Aβ amyloid and RIPK1/RIPK3. **a**. Solid-state NMR structure of the human RIPK1/RIPK3 necrosome was shown in the left panel (PDB 5V7Z), and the crystal structure of the RIPK3 core region was shown in the right panel. **b**. Structural model for Aβ40 fibril with two-fold symmetry was shown in the left panel (PDB 2LMO, by NMR), and the one with three-fold symmetry was shown in the right panel (PDB 2LMP, by NMR). Key residues LMVG are colored in red. **c**. Protein structure alignment of Aβ amyloid (PDB 2LMO and PDB 2LMP, green; PDB 6W0O, cyan) with the tetrapeptide VQVG (red). **d-e**. The polymerization of wild-type, mutated (LMVG/AAAA), and truncated (△LMVG) Aβ40 (**d**) or Aβ42 (**e**) were monitored by the thioflavin T (ThT) fluorescence. Aβ40 (20 μM) or Aβ42 (20 μM) was incubated with 20 μM ThT at 25 °C. All data are representative of three independent experiments. Related to Figure 1.

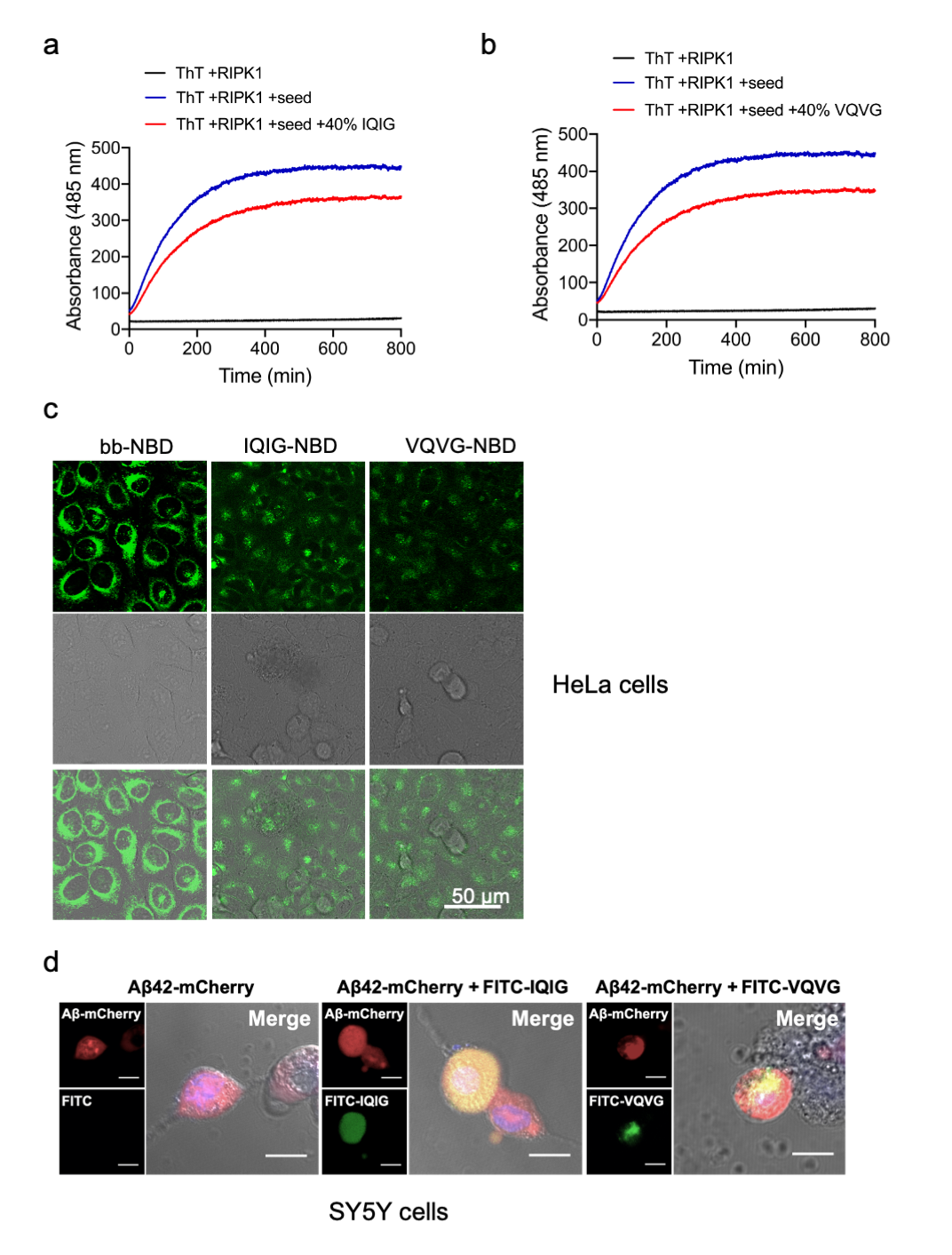

**Figure S2**. Peptides IQIG and VQVG blocked RIPK1 polymerization and decreased TNFα-induced necroptotic cell death. **a-b**. ThT assays for the inhibition of RIPK1 (residues 496-583) polymerization by IQIG (**a**) and VQVG (**b**). RIPK1 (4 μM) was incubated with 10% seed (RIPK1, residues 496-583, polymer), 20 μM ThT, and the corresponding ratio of IQIG or VQVG peptides at 25 °C. **c**. HeLa cells were treated with 2 μM bb-NBD, 5 μM IQIG-NBD, and 5 μM VQVG-NBD, respectively, for 24h before observation by the confocal microscopy. Scale bars, 50 μm. **d**. SY5Y cells were transfected with mCherry-tagged Aβ42. After 24 h, the transfected cells were treated with or without 10 μM FITC-labeled tetrapeptides (IQIG or VQVG) for 6 h. Cellular colocalization between the tetrapeptides and Aβ42 was observed by the confocal microscopy. Cells were fixed and stained with DAPI. Scale bars, 10 μm. FITC: fluorescein isothiocyanate. All data are representative of three independent experiments. Related to Figures 2 and 3.

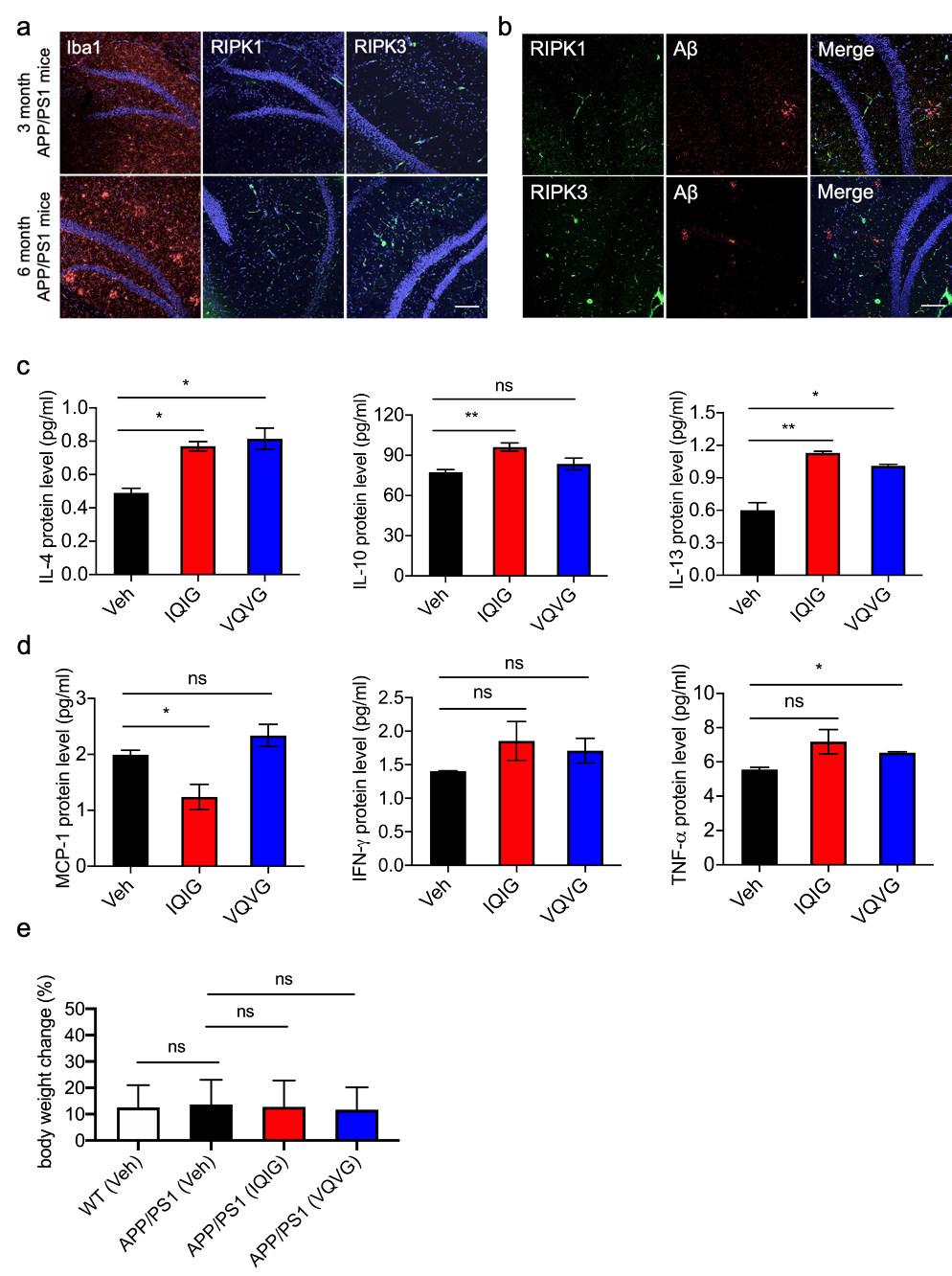

**Figure S3**. IQIG and VQVG reduced the inflammatory response in the APP/PS1 mice. **a**. Representative images of Iba1, RIPK1, and RIPK3 staining in the hippocampus of the APP/PS1 mice (3 and 6 months old). Scale bar: 200 μm. **b**. Representative images showed the distributions of RIPK1 or RIPK3 and Aβ in hippocampal regions of the six-month-old APP/PS1 mice. Scale bar: 200 μm. **c-d**. Cytokine concentrations were measured by the ELISA method in hippocampal lysates from the vehicle-injected, IQIG-injected, and VQVG-injected APP/PS1 mice. **e**. % weight change before and after intravenous administration. The error bars represented the standard deviations (SD). One-way analysis of variance was performed (* *p* < 0.05 and ** *p* < 0.01). All data are representative of three independent experiments. Related to Figure 5.

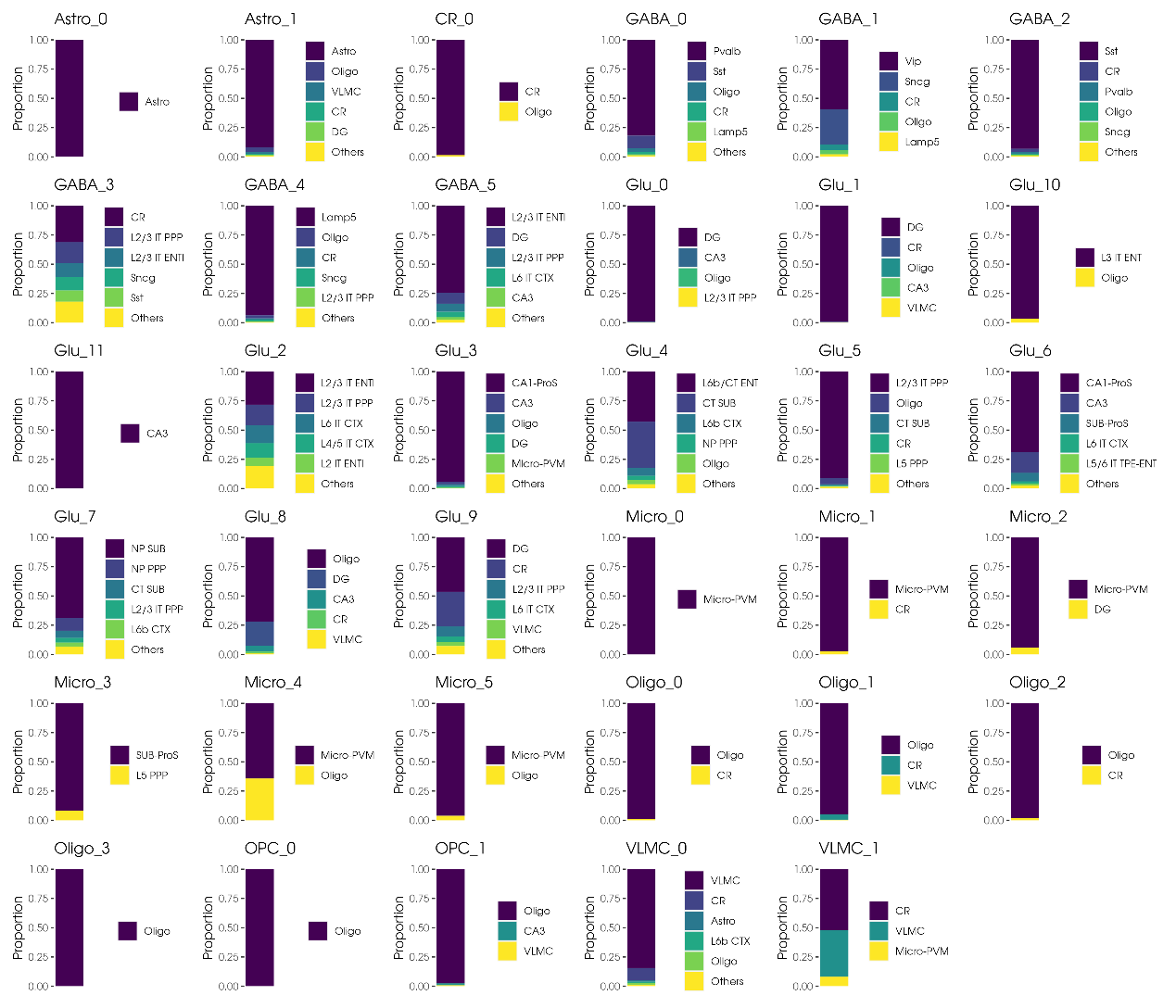

**Figure S4**. Cluster annotation by Seurat. Bar plot showing the predicted cell component of each cluster. Related to Figure 6B, Table S1, Table S4.

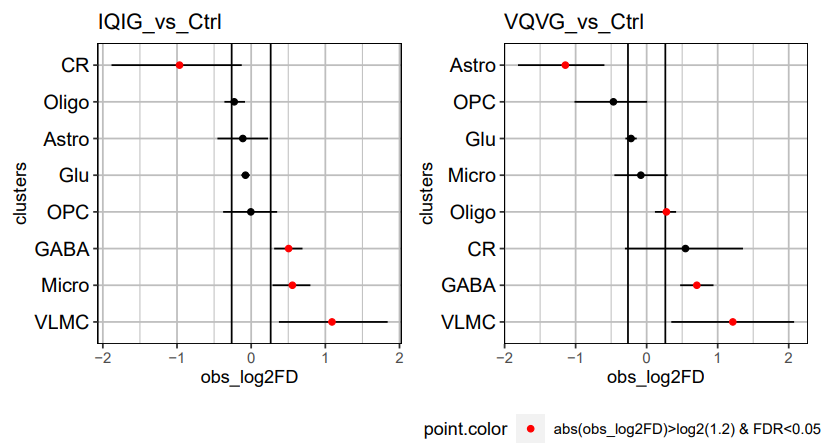

**Figure S5**. Proportion test of main cell type. Related to Figure 6E.

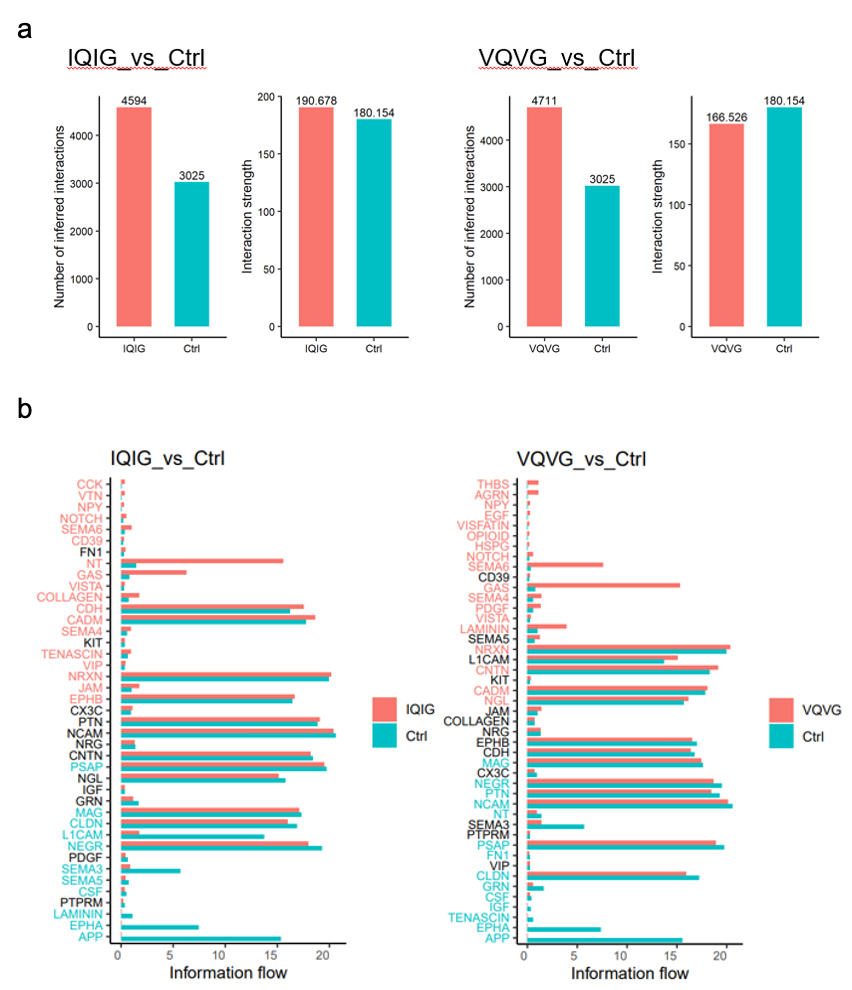

**Figure S6**. Cell-cell communication analysis. **a**. Comparison of total cell-cell communication. The number on each bar represents the total number/strength of inferred cell-cell interactions in each group. **b**. Depiction of context-specific cell-cell communication pathways. While similar to Figure 6G, this figure emphasizes the absolute information flow within each signaling pathway. Related to Figure 6G.

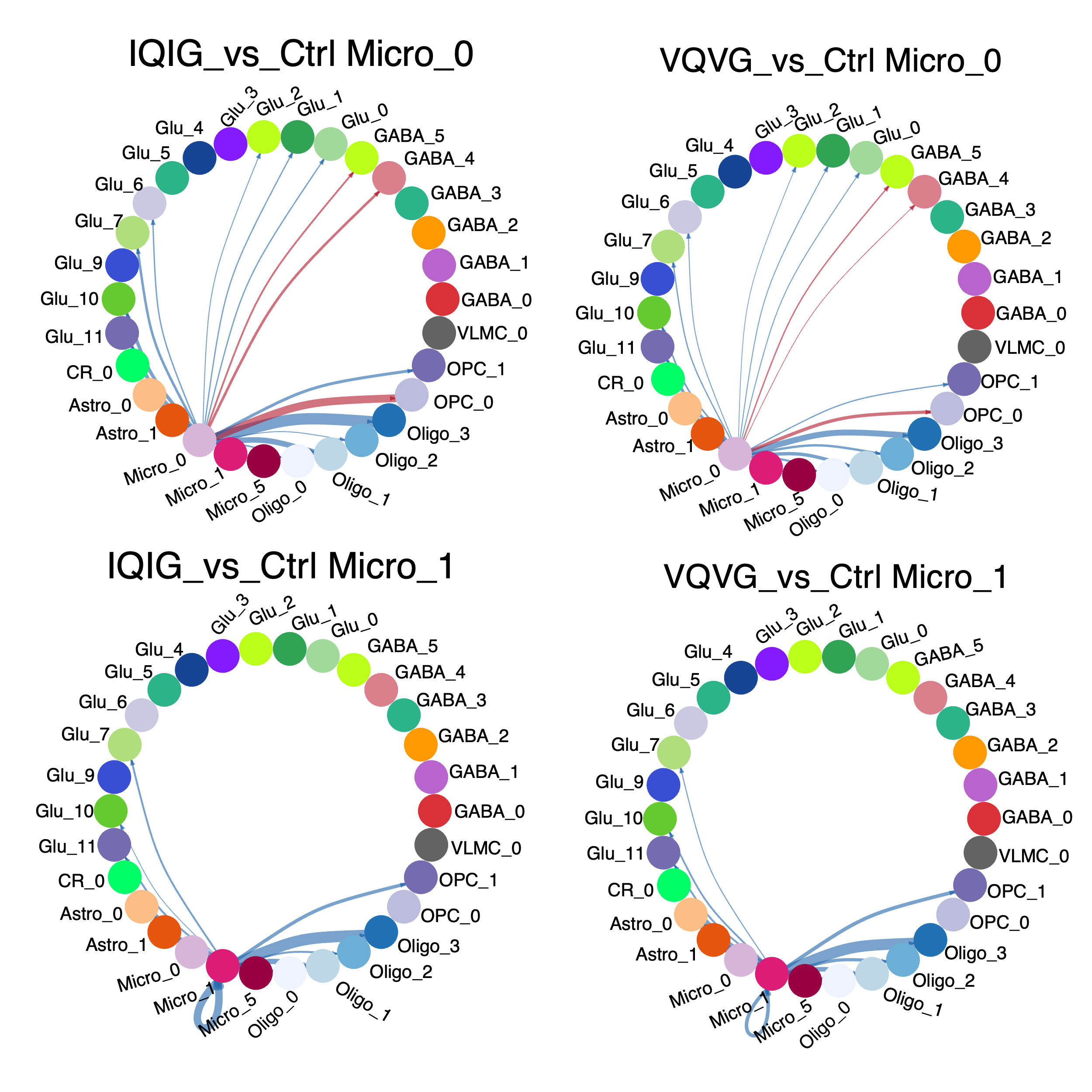

**Figure S7**. The consistent changing pattern of cell-cell communication between Micro_0, Micro_1, and other cell types in two contrasts. This plot shows the consistent change of cell-cell communication between microglia and other cell types in two contrasts (i.e., IQIG-treated vs. vehicle-treated groups (Ctrl), and VQVG-treated vs. vehicle-treated groups). The red (blue) lines inside the circle indicate that the information flow strength is stronger (weaker) in the treatment group than in the control group. Some cell types have less than 10 cells (For example, Micro_1 in VQVG), and their signal strength was set to zero. The line width represents the total signal strength of each cell type. Detailed statistics can be found in Table S4.

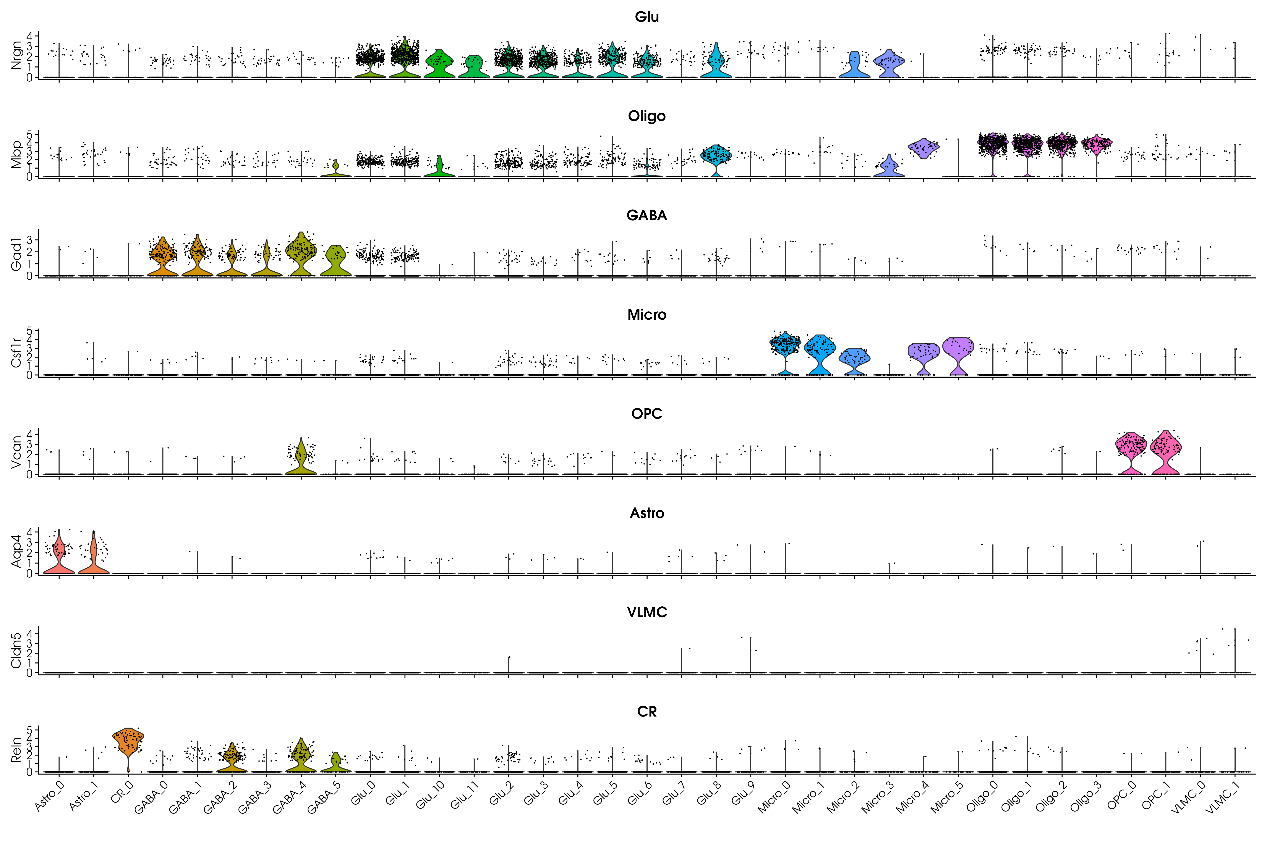

**Figure S8**. Violin plot for all cell type markers to distinguish major cell types.

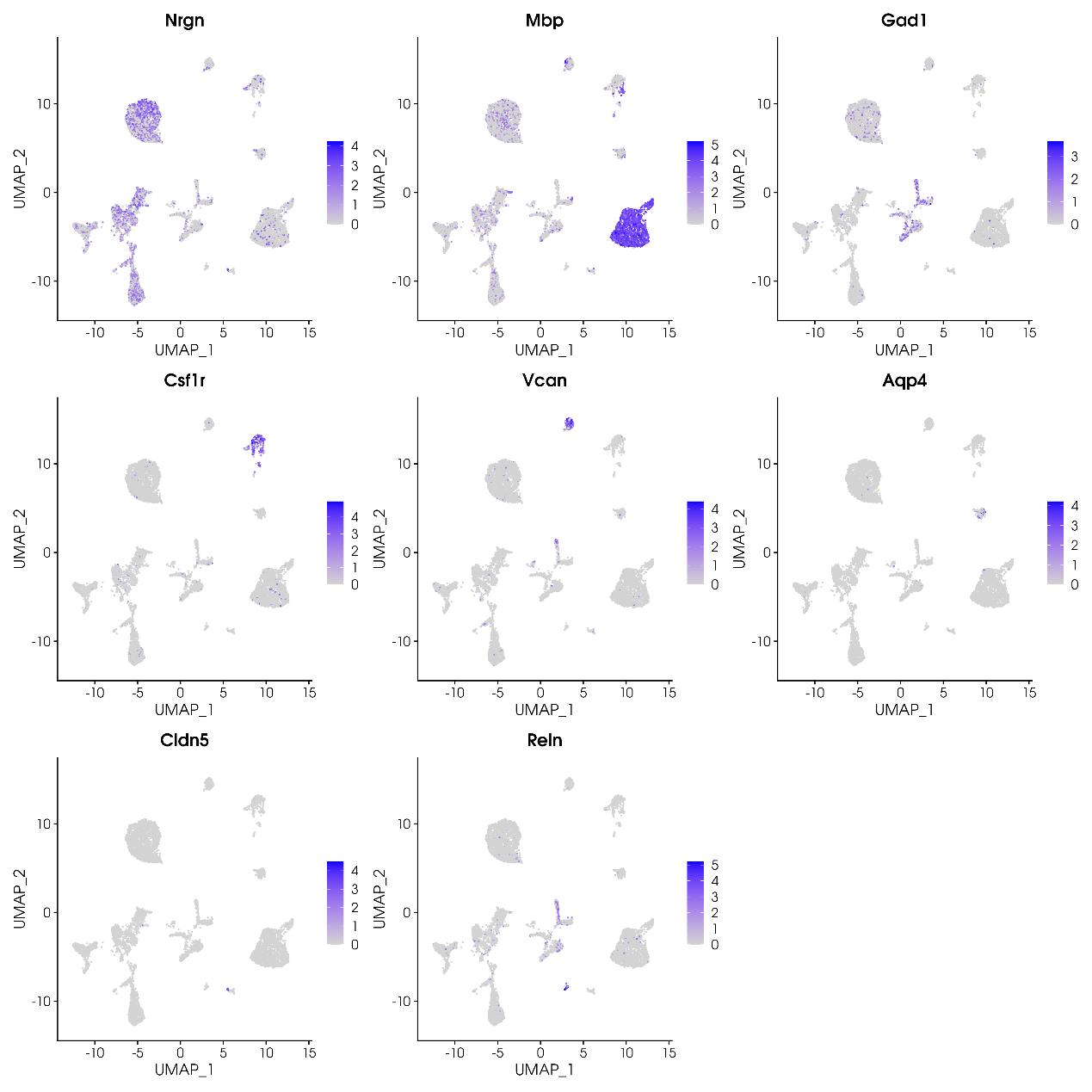

**Figure S9**. Feature plot for all cell type markers to distinguish major cell types.

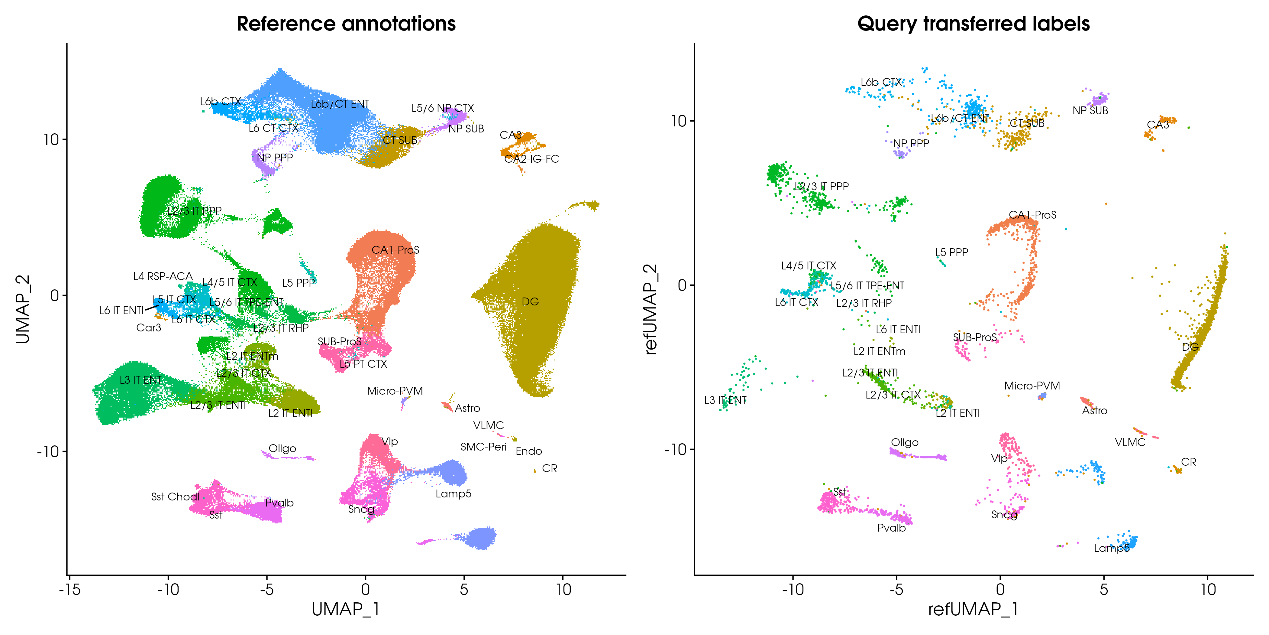

**Figure S10**. Seurat MapQuery mapping plot. The left panel is the reference UMAP, which comes from the hippocampus part of the mouse Whole Cortex, and the Hippocampus 10X reference UMAP structure from the Allen Brain Atlases ^5^. The right UMAP plot represents the projection of our scRNA-seq data to the reference UMAP structure through the MapQuery function of Seurat package.

**SUPPLEMENTARY TABLE LEGENDS**

**Table S1**. Information of each cluster. Including predicted cell identity, color code, cell count, cell proportion, and cell proportion test result between IQIG_vs_Ctrl and VQVG_vs_Ctrl.

**Table S2**. Consistent differential expressed genes between IQIG_vs_Ctrl and VQVG_vs_Ctrl.

**Table S3**. The APP signal pathway between different cell types.

**Table S4**. Cell-cell communication between Micro_0, Micro_1, and other cell types.

**Table S5**. Seurat annotation detail.

**Table S6**. Cell-cell communication between all cell types.
